# Translation initiation or elongation inhibition triggers contrasting effects on *Caenorhabditis elegans* survival during pathogen infection

**DOI:** 10.1101/2024.01.15.575653

**Authors:** Annesha Ghosh, Jogender Singh

## Abstract

Diverse microbial pathogens are known to attenuate host protein synthesis. Consequently, the host mounts a defense response against protein translation inhibition, leading to increased transcript levels of immune genes. The seemingly paradoxical upregulation of immune gene transcripts in response to blocked protein synthesis suggests that the defense mechanism against translation inhibition may not universally benefit host survival. However, a comprehensive assessment of host survival on pathogens upon blockage of different stages of protein synthesis is currently lacking. Here, we investigate the impact of knockdown of various translation initiation and elongation factors on the survival of *Caenorhabditis elegans* exposed to *Pseudomonas aeruginosa*. Intriguingly, we observe opposing effects on *C. elegans* survival depending on whether translation initiation or elongation is inhibited. While translation initiation inhibition enhances survival, elongation inhibition decreases it. Transcriptomic studies reveal that translation initiation inhibition activates a bZIP transcription factor ZIP-2-dependent innate immune response that protects *C. elegans* from *P. aeruginosa* infection. In contrast, inhibiting translation elongation triggers both ZIP-2-dependent and ZIP-2-independent immune responses that, while effective in clearing the infection, are detrimental to the host. Thus, our findings reveal the opposing roles of translation initiation and elongation inhibition in *C. elegans* survival during *P. aeruginosa* infection, highlighting distinct transcriptional reprogramming that may underlie these differences.

## Introduction

Eukaryotic mRNA translation is a highly regulated and energy-consuming process involved in synthesizing proteins from mRNAs (Buttgereit and Brandt, 1995; Jackson et al., 2010). The translation mechanism is critically governed by a combination of initiation, elongation, and release factor polypeptides (Gebauer and Hentze, 2004). Beyond its role in protein synthesis, translation regulates various biological functions such as proteostasis, integrated stress response, neuronal activity, germline apoptosis, and lifespan (Blazie et al., 2021; Clay et al., 2023; Contreras et al., 2008; Curran and Ruvkun, 2007; Gonskikh and Polacek, 2017; Hansen et al., 2007a; Pan et al., 2007; Starck et al., 2016). Different stresses, such as heat shock, nutrient stress, hypoxia, and DNA damage, lead to a general shutdown and reprogramming of translation (Singh, 2020; Spriggs et al., 2010). The integrated stress response triggered by diverse stress signals converges on the global attenuation of translation (Pakos-Zebrucka et al., 2016). A reduced translation rate is essential for survival under stressful conditions (Hansen et al., 2007a; Howard et al., 2016; Rogers et al., 2011), and both translation rate and fidelity play a crucial role in regulating lifespan across various species. Enhanced fidelity of protein synthesis has been linked to increased lifespan (Martinez-Miguel et al., 2021; Xie et al., 2019). Likewise, a reduction in translation rates in otherwise healthy organisms results in an extended lifespan (Hansen et al., 2007a; Pan et al., 2007).

A multitude of polypeptide factors regulate translation initiation and elongation steps (Gebauer and Hentze, 2004; Jackson et al., 2010). Inhibition of translation initiation or elongation can result in different physiological responses (Clay et al., 2023; Knight et al., 2020). For example, inhibiting translation initiation protects against heat and age-associated protein aggregation but does not shield against proteotoxicity caused by proteasome dysfunction. Conversely, inhibiting translation elongation protects against heat and proteasome dysfunction but does not guard against age-associated protein aggregation (Clay et al., 2023). Interestingly, inhibiting both translation initiation and elongation increases the lifespan of the roundworm *Caenorhabditis elegans* (Li et al., 2011; Statzer et al., 2022). Therefore, understanding the role of different translation stages in various physiological processes is important.

Several microbial toxins block host translation (Mohr and Sonenberg, 2012; Remick et al., 2023), prompting organisms to evolve mechanisms that sense perturbations in protein translation and activate immune responses (Chakrabarti et al., 2012; Fontana et al., 2011; Mohr and Sonenberg, 2012). In *C. elegans*, translation inhibition by the bacterial pathogen *Pseudomonas aeruginosa* results in the increased expression of immune genes (Dunbar et al., 2012; Govindan et al., 2015; McEwan et al., 2012). Even in the absence of the pathogen, translation inhibition results in the activation of immune and xenobiotic responses (Dunbar et al., 2012; Govindan et al., 2015; McEwan et al., 2012; Melo and Ruvkun, 2012). Multiple MAP kinases and the bZIP transcription factor ZIP-2 are known to regulate the surveillance immunity activated by translation inhibition (Dunbar et al., 2012; Govindan et al., 2015). Notably, ZIP-2-mediated immune response activation is triggered only by inhibition of translation elongation, not initiation (Dunbar et al., 2012). While the activation of the immune response at the mRNA levels has been monitored, the effects of translation inhibition on the survival of the host against pathogens have not been studied. Increased mRNA levels of immunity genes upon translation inhibition may not necessarily lead to increased production of their polypeptides due to blocked protein synthesis. Thus, a comprehensive analysis of host survival on pathogens upon inhibition of different translation stages is warranted.

In this study, we conducted a comprehensive analysis of the effects of knockdown of different translation initiation and elongation factors on the survival of *C. elegans* on *P. aeruginosa*. Interestingly, we observed that while inhibiting initiation factors either improved or did not change survival, inhibiting elongation factors reduced the survival of *C. elegans* on *P. aeruginosa*. Inhibition of initiation factors activated bZIP transcription factor ZIP-2-dependent protective immune responses. In contrast, inhibition of elongation factors triggered ZIP-2-dependent and ZIP-2-independent immune responses. While these aberrant immune responses helped clear the bacterial infection, they also proved detrimental to the survival of the host. Our findings reveal opposing roles for translation initiation and elongation in the survival of *C. elegans* during *P. aeruginosa* infection and provide insight into the potential molecular mechanisms underlying these differences.

## Results

### Inhibition of translation initiation or elongation leads to opposing outcomes on *C. elegans* survival during *P. aeruginosa* infection

Previous studies have shown that translation inhibition can impact the germline and development of various organisms (Cattie et al., 2016; Contreras et al., 2008; Johnstone and Lasko, 2004; Liu et al., 2007; Long et al., 2002). Therefore, before investigating the impact of inhibiting different translation factors on *C. elegans* survival on a pathogen, we first elucidated how their inhibition affected the fertility and development of *C. elegans*. We systematically knocked down the 47 initiation and elongation factors available in the Ahringer RNA interference (RNAi) library. Among the 47 translation factors tested, the knockdown of 3 resulted in sterility, while the knockdown of 23 slowed the development of *C. elegans* (Table S1). Because sterility modulates survival on pathogens (Miyata et al., 2008), the translation factors whose knockdown led to sterility were excluded from further analysis. For subsequent experiments, we transferred the worms to RNAi plates after the L3 stage for those translation factors whose knockdown slowed the development.

Next, we studied how the knockdown of different translation factors affected the survival of *C. elegans* on *P. aeruginosa* PA14. We observed that the knockdown of different translation factors resulted in an improved (Fig. 1A and B), unaltered (Fig. 1C and D), or reduced (Fig. 1E and F) survival of *C. elegans* compared to the empty vector control on *P. aeruginosa* (Fig. S1; Fig. S2; Table S2). Notably, while the knockdown of translation initiation factors resulted in either increased or unaltered survival, the knockdown of translation elongation factors resulted in reduced survival (Fig. 1G; Fig. S1; Fig. S2; Table S2). Thus, these studies demonstrated that inhibiting translation initiation or elongation led to opposing outcomes on *C. elegans* survival during *P. aeruginosa* infection.

**Figure 1.**
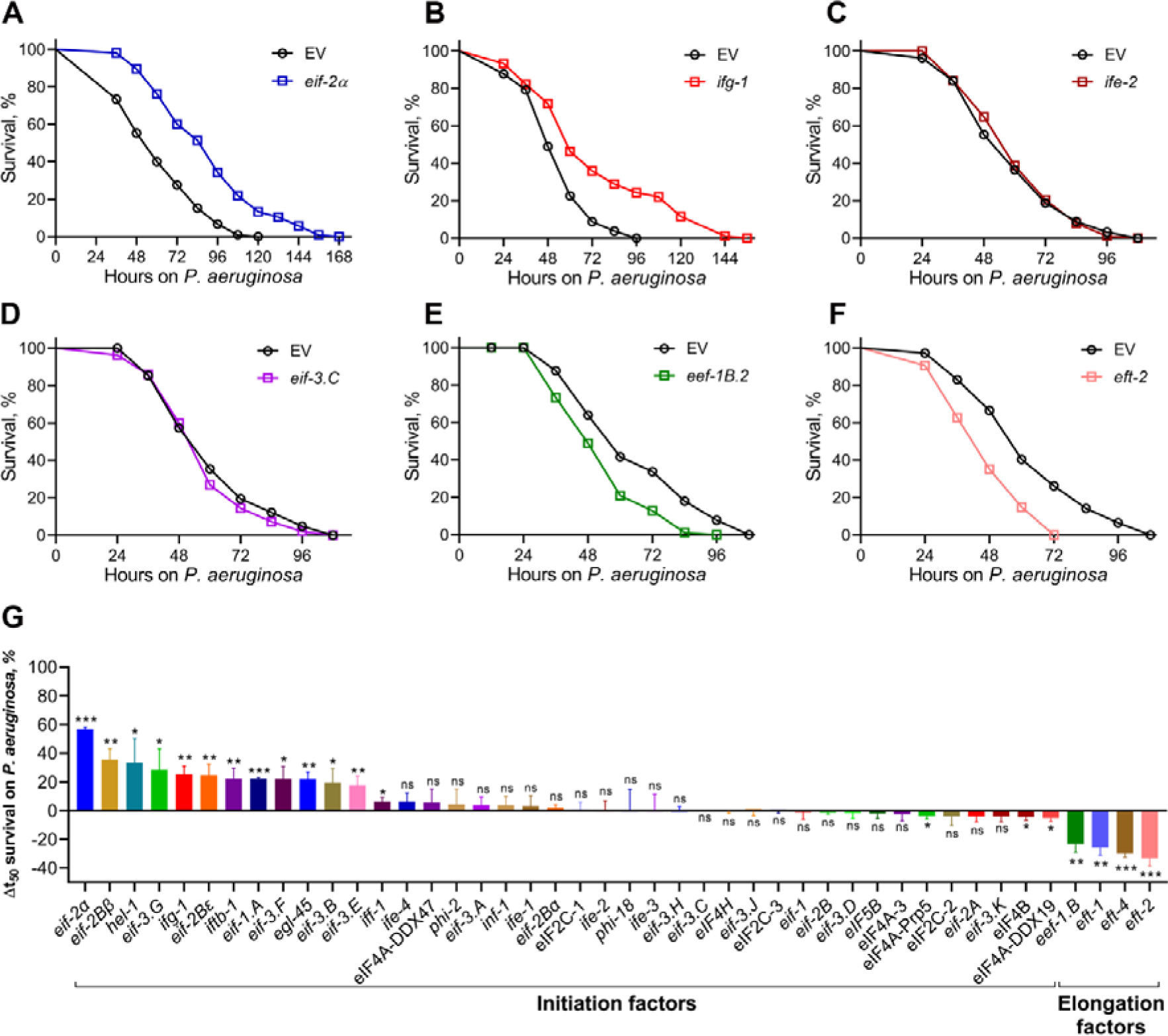
Inhibition of translation initiation or elongation leads to opposing outcomes on *C. elegans* survival during *P. aeruginosa* infection. (A)-(F) Representative survival plots of N2 animals on *P. aeruginosa* PA14 at 25°C after treatment with *eif-2*α (A), *ifg-1* (B), *ife-*2 (C), *eif-3.C* (D), *eef-1B.2* (E), and *eft-2* (F) RNAi, along with the empty vector (EV) control. *P* values for different gene knockdowns compared to the corresponding EV control are as follows: *P*<0.001 for *eif-2*α, *ifg-1*, *eef-1B.2*, and *eft-2*. Survival curves for *ife-2* and *eif-3.C* RNAi compared to EV control are nonsignificant. (G) The percent mean survival change on *P. aeruginosa* upon the knockdown of different initiation and elongation factors compared to the EV control. \*\*\**P* < 0.001, \*\**P* < 0.01, and \**P* < 0.05 via the *t* test. ns, nonsignificant. Data represent the mean and standard deviation from three independent experiments. The triplicate data is available in Table S2.

### Blocking translation initiation enhances *C. elegans* survival on *P. aeruginosa* via the transcription factor ZIP-2

To unravel the immunity pathways contributing to the improved survival of *C. elegans* on *P. aeruginosa* following the inhibition of translation initiation factors, we examined the survival of mutants representing different innate immune pathways. Inhibition of the representative translation initiation factors, *eif-2*α (ortholog of human EIF2S1, eukaryotic translation initiation factor 2 subunit alpha) and *ifg-1* (ortholog of human EIF4G3, eukaryotic translation initiation factor 4 gamma 3), in mutants of various innate immune pathways, including the NSY-1/SEK-1/PMK-1 (Kim et al., 2002), the TGF-β/DBL-1 (Mallo et al., 2002), and the nuclear hormone receptor NHR-8 (Otarigho and Aballay, 2020; Peterson et al., 2022), resulted in enhanced survival compared to control RNAi (Fig. 2A-E; Fig. S3A-E). However, the mutants of the bZIP transcription factor ZIP-2 (Estes et al., 2010) did not exhibit increased survival on *P. aeruginosa* upon the knockdown of *eif-2*α and *ifg-1* (Fig. 2F; Fig. S3F), suggesting that the enhanced survival upon inhibiting translation initiation factors is mediated by ZIP-2.

**Figure 2.**
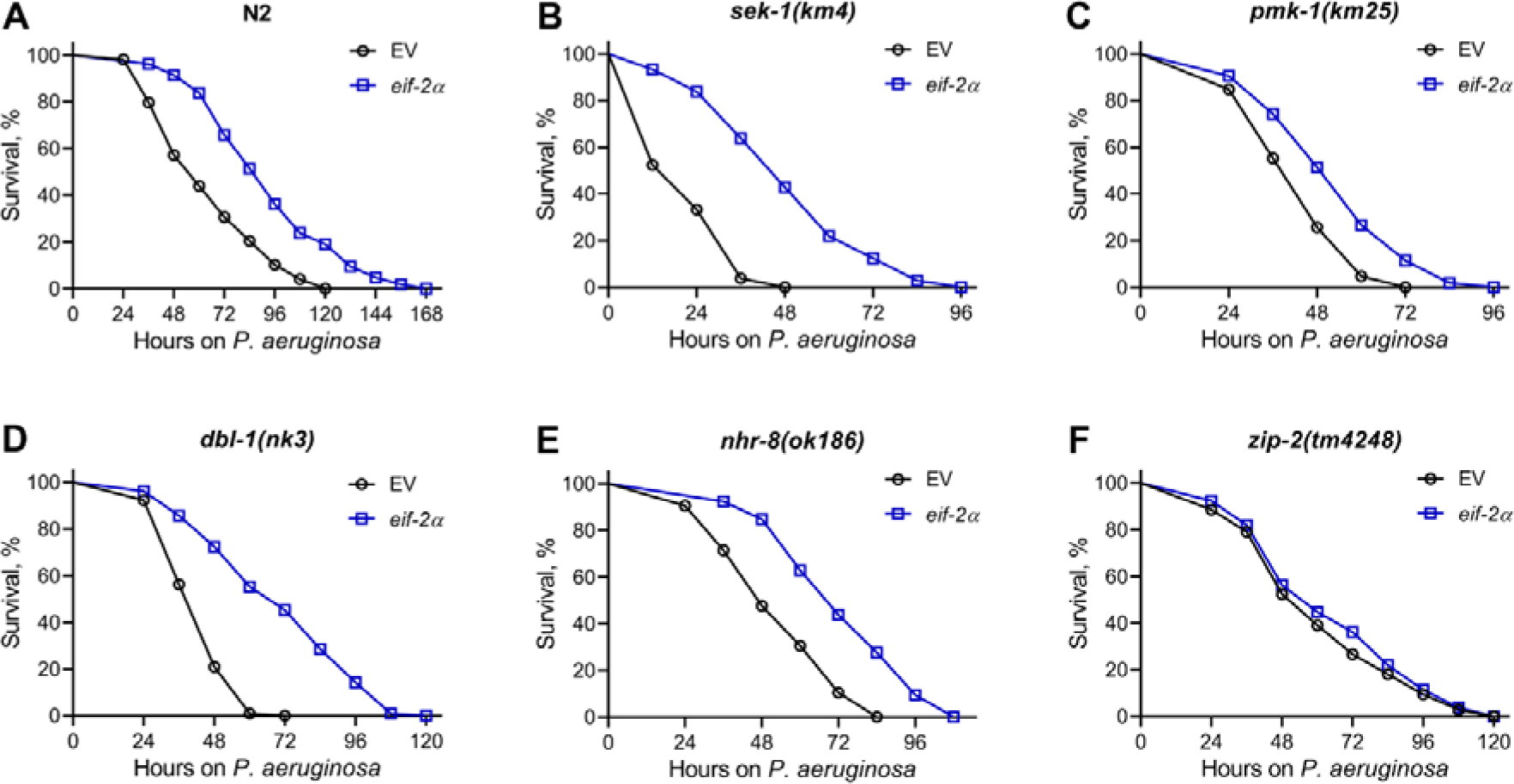
Knockdown of *eif-2*α increases *C. elegans* survival on *P. aeruginosa* via transcription factor ZIP-2. Representative survival plots of N2 (A), *sek-1(km4)* (B), *pmk-1(km25)* (C), *dbl-1(nk3)* (D), *nhr-8(ok186)* (E), and *zip-2(tm4248)* (F) animals on *P. aeruginosa* PA14 at 25°C after treatment with the empty vector (EV) control and *eif-2*α RNAi. *P*<0.001 for *eif-2*α RNAi compared to EV control for N2, *sek-1(km4)*, *pmk-1(km25)*, *dbl-1(nk3)*, and *nhr-8(ok186)*. Survival curves for *eif-2*α RNAi compared to EV control for *zip-2(tm4248)* are nonsignificant.

Since the knockdown of *eif-2*α and *ifg-1* did not show a significant difference in the survival rate of *zip-2(tm4248)* animals on *P. aeruginosa* compared to control RNAi animals, it is important to test the sensitivity of *zip-2(tm4248)* animals to RNAi. Therefore, we knocked down *act-5* and *bli-3* genes in N2 wild-type, *zip-2(tm4248)*, and RNAi-defective animals *sid-1(qt9)*. The knockdown of *act-5* and *bli-3* is known to result in development arrest and a blistered cuticle, respectively (Gahlot and Singh, 2024; Simmer et al., 2003; Whangbo et al., 2017). We observed that N2 and *zip-2(tm4248)* animals were arrested at the L1 stage upon *act-5* RNAi and developed multiple blisters across the larval body upon *bli-3* RNAi (Fig. S4). In contrast, the RNAi-defective *sid-1(qt9)* animals did not display these phenotypes and developed into healthy adults on *act-5* and *bli-3* RNAi (Fig. S4). These results indicated that *zip-2(tm4248)* animals have sensitivity to RNAi similar to N2 animals, and the enhanced survival on *P. aeruginosa* upon inhibiting translation initiation factors is likely mediated by ZIP-2.

### Inhibition of translation initiation activates a ZIP-2-dependent immune response

Previous studies have shown that the inhibition of translation initiation and elongation results in the increased expression of innate immune and xenobiotic response genes (Dunbar et al., 2012; Govindan et al., 2015). While immune gene induction upon translation elongation inhibition was found to be ZIP-2 dependent, translation initiation inhibition was initially thought to induce immune genes independently of ZIP-2 (Dunbar et al., 2012). However, given that the enhanced survival of *C. elegans* on *P. aeruginosa* upon translation initiation inhibition was entirely dependent on ZIP-2, we hypothesized that this inhibition might indeed trigger ZIP-2-dependent immune responses. To explore this, we conducted RNA sequencing to examine transcriptional changes in N2 and *zip-2(tm4248)* animals following *eif-2*α knockdown (Tables S3 and S4). Gene Ontology (GO) analysis of upregulated genes in N2 animals revealed enrichment of innate immune response (Fig. 3A) and structural constituents of the cuticle, including collagen genes (Fig. 3B). In contrast, downregulated genes in N2 animals were enriched for processes such as cell cycle, cell division, and translation (Fig. S5A and B). This included significant downregulation of *eif-2*α mRNA (Table S3), confirming efficient RNAi-mediated suppression of its transcript levels. The mRNA levels of several other translation initiation factor genes were also downregulated, suggesting a coordinated downregulation of multiple initiation factors upon inhibition of *eif-2*α expression.

**Figure 3.**
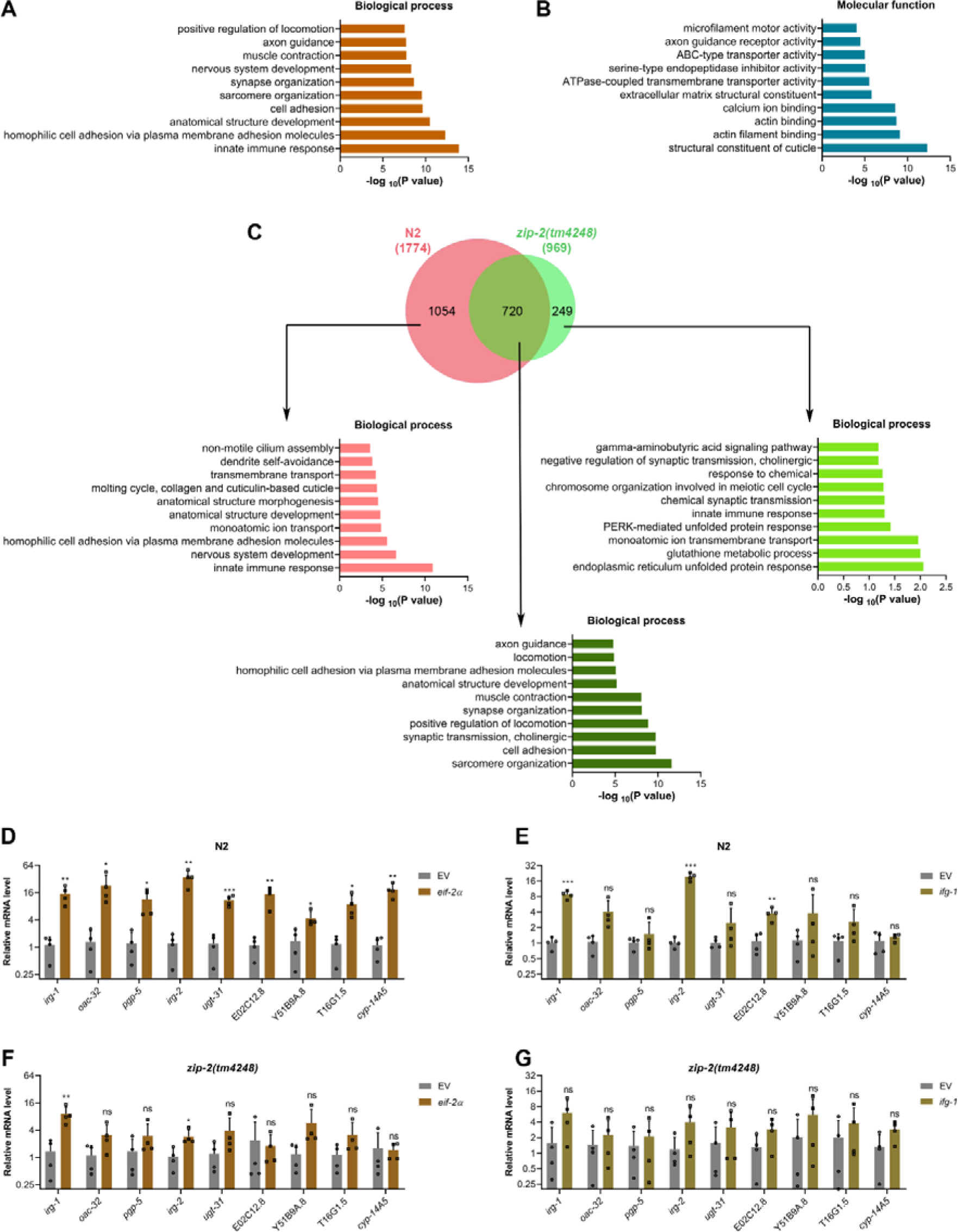
Inhibition of translation initiation activates a ZIP-2-dependent immune response. (A)-(B) Gene Ontology (GO) enrichment analysis of upregulated genes upon *eif-2*α knockdown in N2 animals for biological processes (A) and molecular functions (B). (C) Venn diagram showing the overlap between genes upregulated upon *eif-2*α knockdown in N2 and *zip-2(tm4248)* animals. The GO analysis for biological processes of unique and common genes is shown. (D) Quantitative reverse transcription-PCR (qRT-PCR) for immune genes expression analysis of N2 animals after treatment with the empty vector (EV) control and *eif-2*α RNAi. (E) qRT-PCR for immune genes expression analysis of N2 animals after treatment with the EV control and *ifg-1* RNAi. (F) qRT-PCR for immune genes expression analysis of *zip-2(tm4248)* animals after treatment with the EV control and *eif-2*α RNAi. (G) qRT-PCR for immune genes expression analysis of *zip-2(tm4248)* animals after treatment with the EV control and *ifg-1* RNAi. For panels (D)-(G), \*\*\**P* < 0.001, \*\**P* < 0.01, and \**P* < 0.05 via the *t* test. ns, nonsignificant. Data represent the mean and standard deviation from four independent experiments.

A comparison of *eif-2*α RNAi-upregulated genes in N2 and *zip-2(tm4248)* animals revealed that the expression of 1054 genes was ZIP-2 dependent (Fig. 3C; Table S5), with a strong enrichment for innate immune responses. Notably, collagen gene upregulation was also ZIP-2 dependent (Fig. S5C). Genes upregulated independently of ZIP-2 did not show enrichment for innate immune responses (Fig. 3C; Fig. S5C). To validate the RNA sequencing data, we used quantitative reverse transcription-PCR (qRT-PCR) to monitor the mRNA levels of selected immune genes that were upregulated in N2 but not in *zip-2(tm4248)*. The *eif-2*α knockdown strongly induced these genes (Fig. 3D), whereas the *ifg-1* knockdown had a weaker effect, possibly explaining the differences in survival rates on *P. aeruginosa* (Fig. 1G). The expression levels of these genes declined substantially in *zip-2(tm4248)* animals upon knockdown of *eif-2*α and *ifg-1*, suggesting that their induction was at least partially ZIP-2 dependent (Fig. 3F and G). Previous studies have shown that translation inhibition activates ZIP-2 signaling by increasing its expression (Dunbar et al., 2012; Reddy et al., 2016). We found that *eif-2*α knockdown led to about 4-fold increase in *zip-2* mRNA levels (Table S3), suggesting that translation initiation inhibition may regulate ZIP-2 signaling at the transcription level. We also tested the expression levels of genes from other immunity pathways and found that those were not induced in N2 and *zip-2(tm4248)* animals upon knockdown of *eif-2*α and *ifg-1* (Fig. S5D-G). Collectively, these results indicated that translation initiation inhibition activates a ZIP-2-dependent immune response, contributing to enhanced survival on *P. aeruginosa*.

The colonization of the *C. elegans* gut by *P. aeruginosa* plays a critical role in infection and survival under slow-killing conditions (Das et al., 2023; Tan et al., 1999). We investigated whether inhibiting translation initiation factors affected gut colonization by *P. aeruginosa*. Knockdown of *ifg-1* and *eif-2*α in N2 animals significantly reduced gut colonization (Fig. 4A and B). To further validate this, we measured colony-forming units (CFU) of *P. aeruginosa* in worms after *ifg-1* and *eif-2*α knockdown. Knockdown of *ifg-1* and *eif-2*α led to a significant decrease in CFU per worm, indicating fewer live bacteria (Fig. 4C). We also evaluated the impact of *ifg-1* and *eif-2*α knockdown on *P. aeruginosa* colonization in different immunity mutants. Inhibition of *ifg-1* and *eif-2*α also reduced *P. aeruginosa* colonization in *sek-1(km4)*, *pmk-1(km25)*, *dbl-1(nk3)*, and *nhr-8(ok186)* animals (Fig. S6). Interestingly, while *ifg-1* knockdown reduced colonization in *zip-2(tm4248)* animals, *eif-2*α knockdown only partially reduced colonization (Fig. 4D-F), indicating that ZIP-2 may play a minor role in regulating gut colonization.

**Figure 4.**
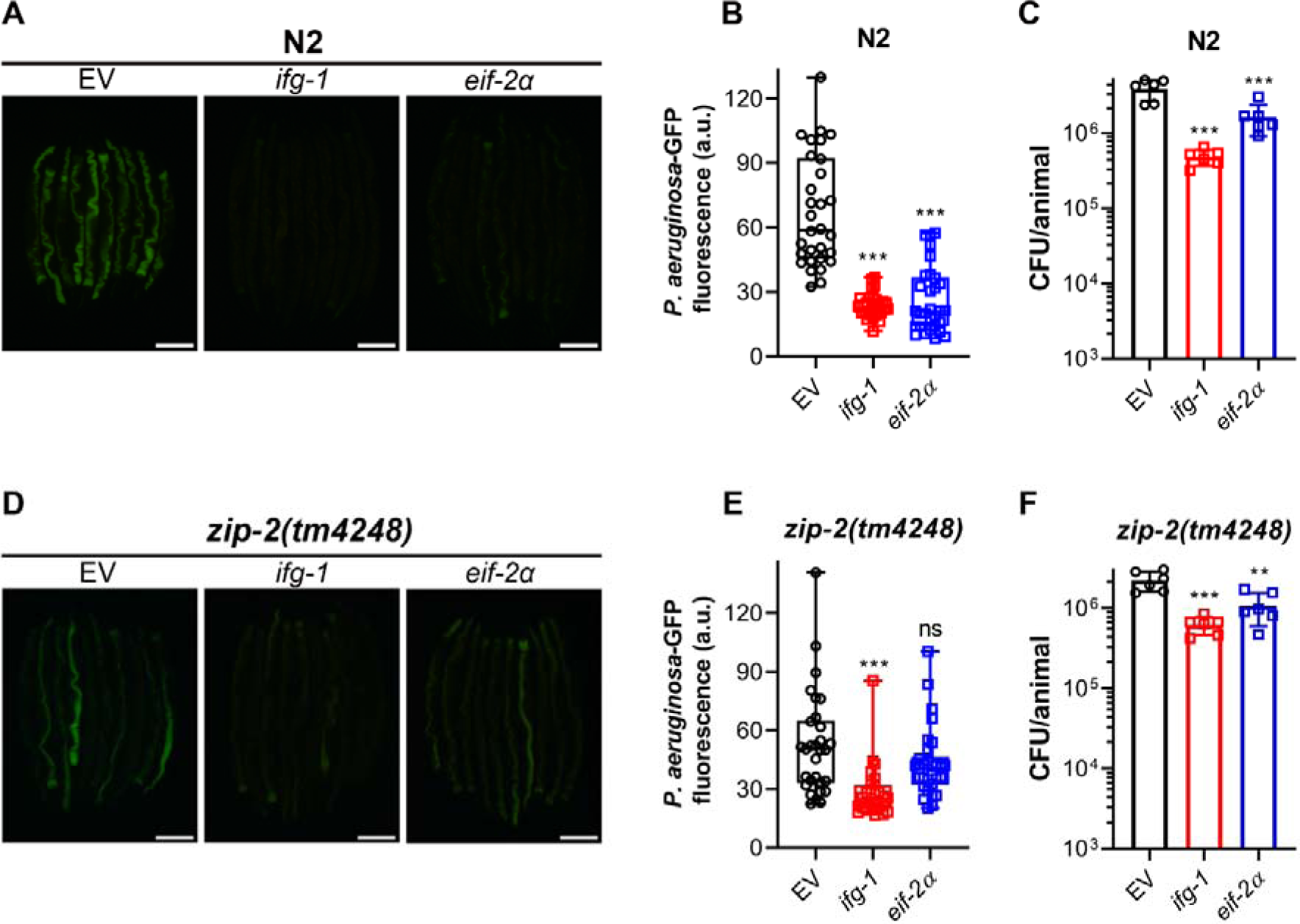
Inhibition of translation initiation reduces *C. elegans* gut colonization with *P. aeruginosa*. (A) Representative fluorescence images of N2 animals incubated on *P. aeruginosa*-GFP for 24 hours at 25°C after treatment with the empty vector (EV) control, *ifg-1*, and *eif-2*α RNAi. Scale bar = 200 μm. (B) Quantification of GFP levels of N2 animals incubated on *P. aeruginosa*-GFP for 24 hours at 25°C after treatment with the EV control, *ifg-1*, and *eif-2*α RNAi. \*\*\**P* < 0.001 via ordinary one-way ANOVA followed by Dunnett’s multiple comparisons test (*n* = 30 worms each). (C) Colony-forming units (CFU) per animal of N2 worms incubated on *P. aeruginosa*-GFP for 24 hours at 25°C after treatment with the EV control, *ifg-1*, and *eif-2*α RNAi. \*\*\**P* < 0.001 via ordinary one-way ANOVA followed by Dunnett’s multiple comparisons test (*n = 6* biological replicates). (D) Representative fluorescence images of *zip-2(tm4248)* animals incubated on *P. aeruginosa*-GFP for 24 hours at 25°C after treatment with the EV control, *ifg-1*, and *eif-2*α RNAi. Scale bar = 200 μm. (E) Quantification of GFP levels of *zip-2(tm4248)* animals incubated on *P. aeruginosa*-GFP for 24 hours at 25°C after treatment with the EV control, *ifg-1*, and *eif-2*α RNAi. \*\*\**P* < 0.001 and ns, nonsignificant via ordinary one-way ANOVA followed by Dunnett’s multiple comparisons test (*n* = 30 worms each). (F) CFU per animal of *zip-2(tm4248)* worms incubated on *P. aeruginosa*-GFP for 24 hours at 25°C after treatment with the EV control, *ifg-1*, and *eif-2*α RNAi. \*\*\**P* < 0.001 and \*\**P* < 0.01 via ordinary one-way ANOVA followed by Dunnett’s multiple comparisons test (*n = 6* biological replicates).

Notably, *ifg-1* knockdown led to lower *P. aeruginosa* colonization levels than *eif-2*α knockdown in N2 and various immunity mutants, including *zip-2(tm4248)*. However, *eif-2*α knockdown resulted in better survival on *P. aeruginosa* compared to *ifg-1* knockdown (Fig. 1G). This suggested that the reduction in *P. aeruginosa* colonization does not correlate with improved survival, indicating that gut colonization may not be a reliable predictor of *C. elegans* survival on *P. aeruginosa*.

### Detrimental effects of translation elongation inhibition on *C. elegans* survival on *P. aeruginosa* diminish in *zip-2(tm4248)* animals

To understand how inhibiting translation elongation reduces the survival of *C. elegans* on *P. aeruginosa*, we examined the effects of knocking down two representative elongation factors: *eef-1B.2* (an ortholog of human EEF1B2, eukaryotic elongation factor 1 beta 2) and *eft-2* (an ortholog of human EEF2, eukaryotic elongation factor 2). We tested the survival of mutants from different innate immune pathways after the knockdown of these elongation factors. While the knockdown of *eef-1B.2* resulted in the reduced survival of *pmk-1(km25)*, *dbl-1(nk3)*, and *nhr-8(ok186)* animals, its knockdown did not affect the survival of *sek-1(km4)* and *zip-2(tm4248)* animals (Fig. 5). Given that SEK-1 and PMK-1 are part of the same signaling pathway (Kim et al., 2002), the reduction in survival observed in *pmk-1(km25)* but not *sek-1(km4)* animals makes it difficult to determine whether the decreased survival due to *eef-1B.2* knockdown is SEK-1 dependent. These differences may arise because, compared to the *pmk-1* knockout, the *sek-1* knockout has more severe effects on survival following infection with *P. aeruginosa* (Meng et al., 2021; Rao et al., 2024). We also studied the survival of these immune pathway mutants following *eft-2* knockdown. Interestingly, in contrast to reducing survival of N2, the knockdown of *eft-2* resulted in improved survival of *sek-1(km4)*, *dbl-1(nk3)*, and *nhr-8(ok186)* animals on *P. aeruginosa* (Fig. 6). There was no effect of *eft-2* knockdown on the survival of *pmk-1(km25)* and *zip-2(tm4248)* animals on *P. aeruginosa* (Fig. 6C and F). It is difficult to assess whether the survival on *P. aeruginosa* upon *eft-2* knockdown is PMK-1 dependent as *sek-1(km4)* animals exhibited increased survival upon *eft-2* knockdown. Nevertheless, the detrimental effects of *eef-1B.2* and *eft-2* knockdown diminished in *zip-2(tm4248)* animals (Fig. 5F; Fig. 6F). Notably, when comparing N2 and all immunity pathway mutants, *zip-2(tm4248)* animals displayed the best survival on *P. aeruginosa* following *eef-1B.2* and *eft-2* knockdown among all the strains (Fig. S7), suggesting that *zip-2(tm4248)* animals had least detrimental effects of *eef-1B.2* and *eft-2* knockdown.

**Figure 5.**
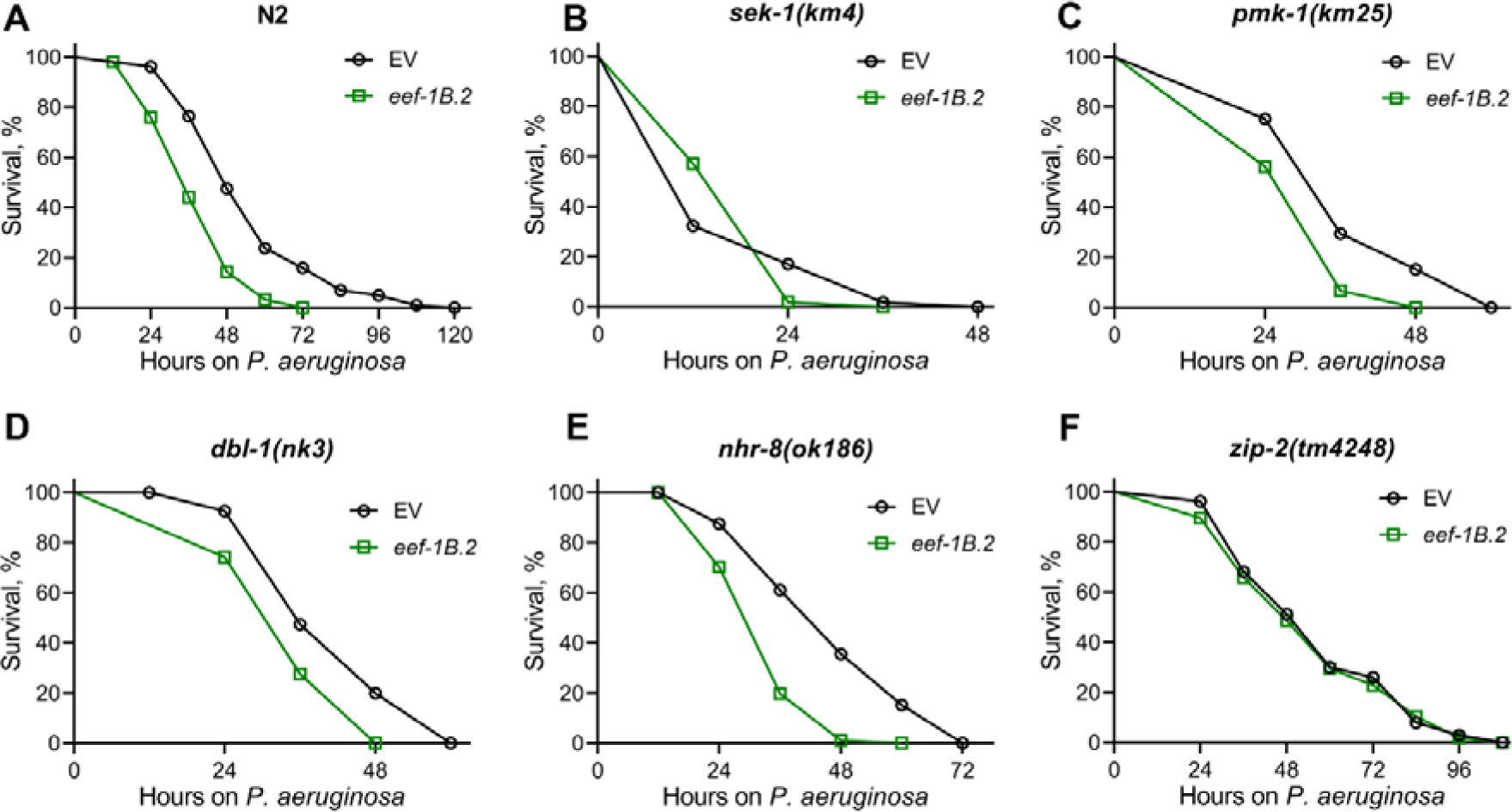
Detrimental effects of knockdown of *eef-1B.2* on *C. elegans* survival on *P. aeruginosa* diminish in *zip-2(tm4248)* animals. Representative survival plots of N2 (A), *sek-1(km4)* (B), *pmk-1(km25)* (C), *dbl-1(nk3)* (D), *nhr-8(ok186)* (E), and *zip-2(tm4248)* (F) animals on *P. aeruginosa* PA14 at 25°C after treatment with the empty vector (EV) control and *eef-1B.2* RNAi. *P*<0.001 for *eef-1B.2* RNAi compared to EV control for N2, *pmk-1(km25)*, *dbl-1(nk3)*, and *nhr-8(ok186)*. Survival curves for *eef-1B.2* RNAi compared to EV control for *sek-1(km4)* and *zip-2(tm4248)* are nonsignificant.

**Figure 6.**
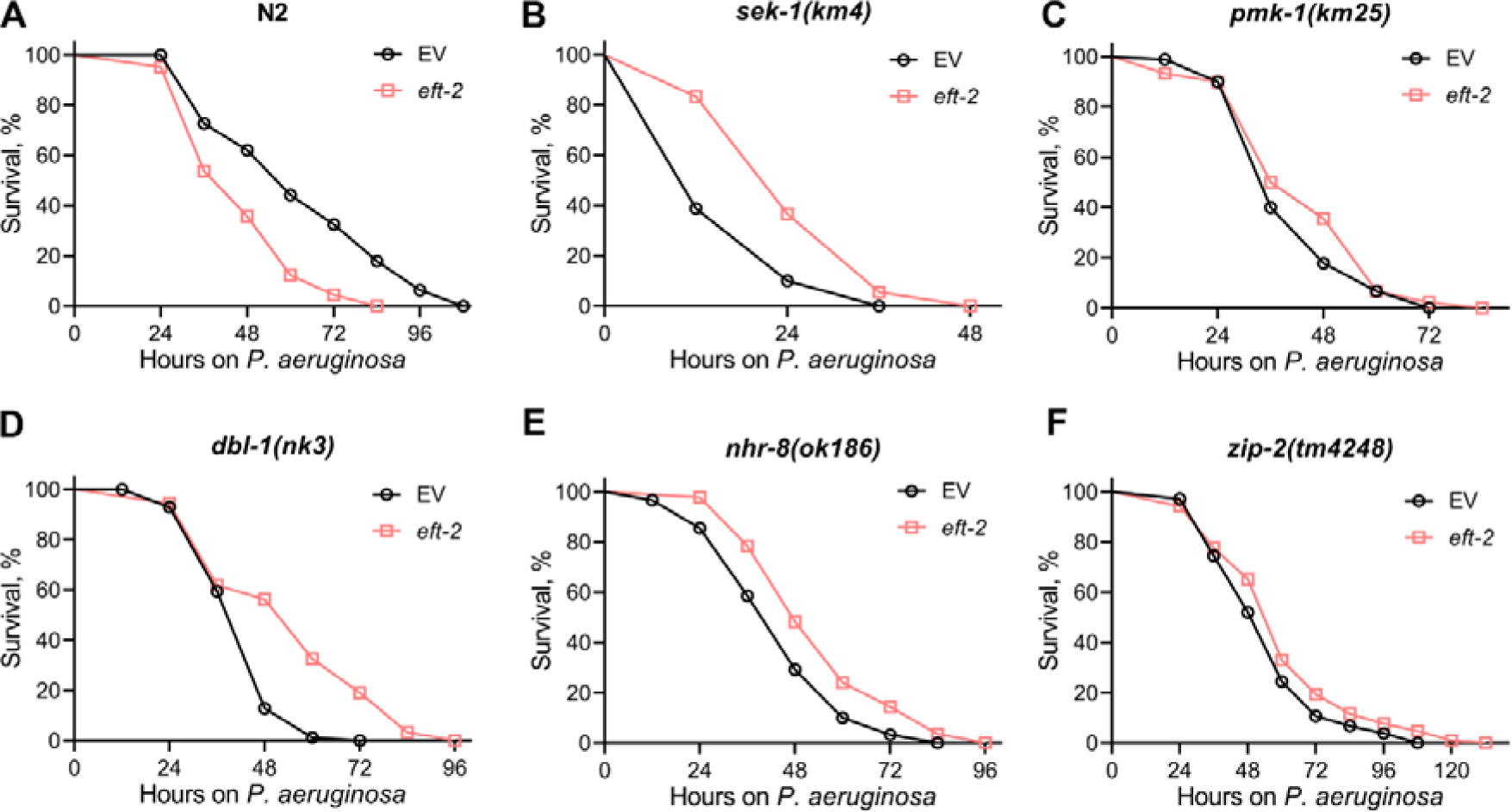
Detrimental effects of knockdown of *eft-2* on *C. elegans* survival on *P. aeruginosa* diminish in *zip-2(tm4248)* animals. Representative survival plots of N2 (A), *sek-1(km4)* (B), *pmk-1(km25)* (C), *dbl-1(nk3)* (D), *nhr-8(ok186)* (E), and *zip-2(tm4248)* (F) animals on *P. aeruginosa* PA14 at 25°C after treatment with the empty vector (EV) control and *eft-2* RNAi. *P*<0.001 for *eft-2* RNAi compared to EV control for N2, *sek-1(km4)*, *dbl-1(nk3)*, and *nhr-8(ok186)*. *P*<0.05 for *eft-2* RNAi compared to EV control for *zip-2(tm4248)*. Survival curves for *eft-2* RNAi compared to EV control for *pmk-1(km25)* are nonsignificant.

### Inhibition of translation elongation activates ZIP-2-dependent and ZIP-2-independent immune responses

To understand the molecular basis of the detrimental effects of inhibiting translation elongation on *C. elegans* survival during *P. aeruginosa* infection, we characterized transcriptomic changes in N2 and *zip-2(tm4248)* animals after *eft-2* knockdown (Tables S6 and S7). GO analysis for biological processes of upregulated genes in N2 animals revealed a strong enrichment for innate immune responses (Fig. 7A). The enrichment for innate immune responses appeared much stronger for *eft-2* knockdown compared to *eif-2*α knockdown (Fig. 3A; Fig. 7A). GO analysis for molecular functions highlighted enrichment for transcription factors and ABC-type transporter (Fig. 7B). Unlike *eif-2*α knockdown, *eft-2* knockdown did not show enrichment for constituents of the cuticle (Fig. 3B; Fig. 7B). GO analysis of downregulated genes in N2 animals indicated enrichment for processes like cell death and D-amino acid oxidase activity (Fig. S8A and B). Additionally, processes related to proteostasis, including protein ubiquitination and endopeptidase activity, were enriched among the downregulated genes. This suggested that the reduced protein synthesis load may lead to the downregulation of genes involved in protein degradation.

**Figure 7.**
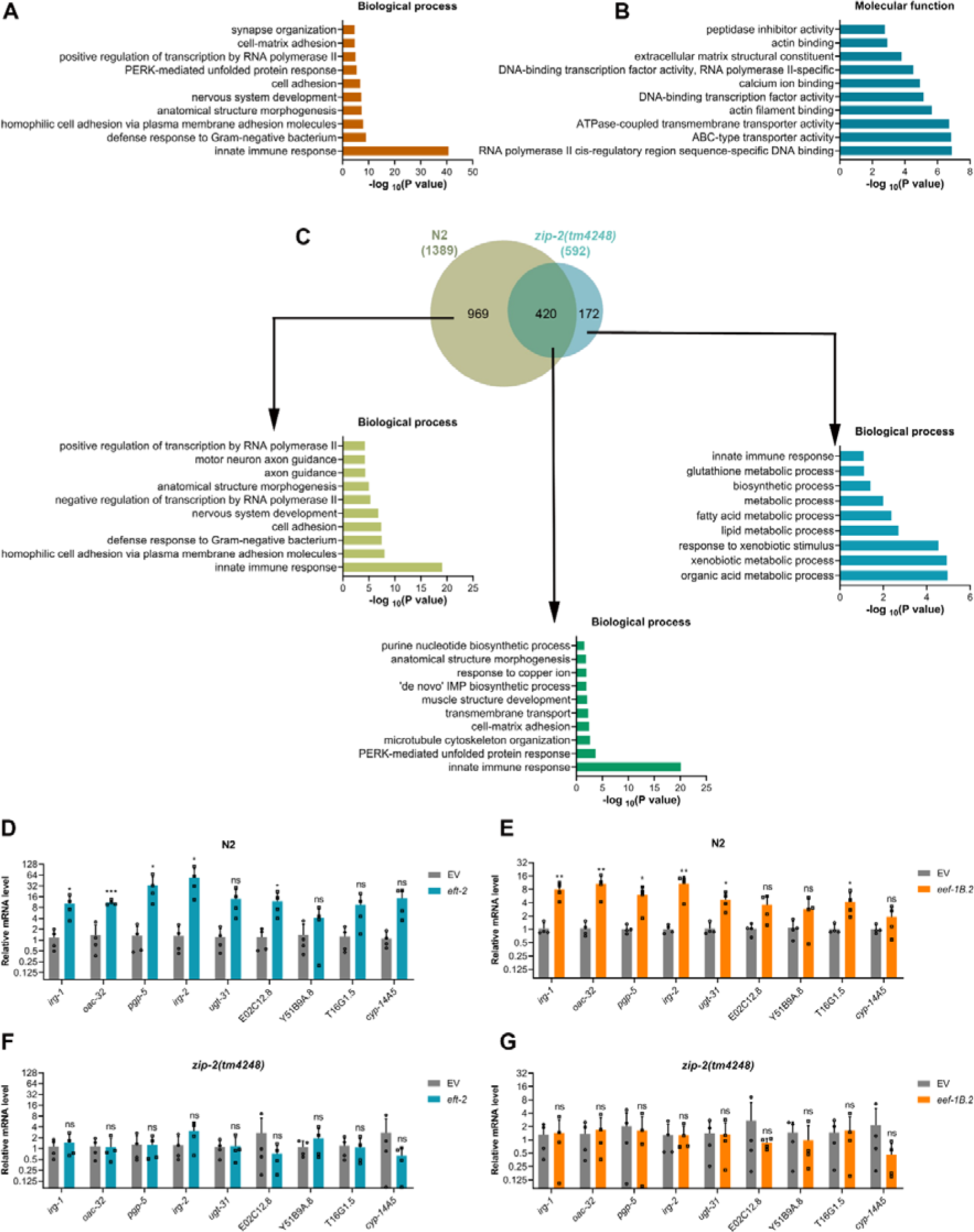
Inhibition of translation elongation activates ZIP-2-dependent and ZIP-2-independent immune responses. (A)-(B) Gene Ontology (GO) enrichment analysis of upregulated genes upon *eft-2* knockdown in N2 animals for biological processes (A) and molecular functions (B). (C) Venn diagram showing the overlap between genes upregulated upon *eft-2* knockdown in N2 and *zip-2(tm4248)* animals. The GO analysis for biological processes of unique and common genes is shown. (D) Quantitative reverse transcription-PCR (qRT-PCR) for immune genes expression analysis of N2 animals after treatment with the empty vector (EV) control and *eft-2* RNAi. (E) qRT-PCR for immune genes expression analysis of N2 animals after treatment with the EV control and *eef-1B.2* RNAi. (F) qRT-PCR for immune genes expression analysis of *zip-2(tm4248)* animals after treatment with the EV control and *eft-2* RNAi. (G) qRT-PCR for immune genes expression analysis of *zip-2(tm4248)* animals after treatment with the EV control and *eef-1B.2* RNAi. For panels (D)-(G), \*\*\**P* < 0.001, \*\**P* < 0.01, and \**P* < 0.05 via the *t* test. ns, nonsignificant. Data represent the mean and standard deviation from four independent experiments.

A comparison of the *eft-2* RNAi upregulated genes in N2 and *zip-2(tm4248)* animals showed that the expression of 969 genes was ZIP-2 dependent (Fig. 7C; Table S8). These ZIP-2-dependent genes were strongly enriched for innate immune response. Interestingly, ZIP-2-independent genes upregulated in N2 also showed a strong enrichment for innate immune responses (Fig. 7C), along with MAP kinase activity (Fig. S8C). To validate the RNA sequencing data, we measured the mRNA levels of ZIP-2-dependent immunity genes by qRT-PCR. Most of the tested genes had significant induction upon the knockdown of *eft-2* and *eef-1B.2* in N2 animals (Fig. 7D and E). This induction was ZIP-2 dependent, as these genes were not induced in *zip-2(tm4248)* animals after *eft-2* and *eef-1B.2* knockdown (Fig. 7F and G). Similar to *eif-2*α knockdown, *eft-2* knockdown also resulted in increased levels of *zip-2* mRNA (Table S6). We also tested the expression levels of genes from other immunity pathways and found that they were not induced in N2 and *zip-2(tm4248)* animals following *eft-2* and *eef-1B.2* knockdown (Fig. S8D-G). Taken together, these results suggested that the inhibition of translation elongation activated ZIP-2-dependent and ZIP-2-independent immune responses.

### ZIP-2 reduces gut colonization by *P. aeruginosa* upon inhibition of translation elongation

Given that inhibiting translation elongation led to reduced survival in N2 animals on *P. aeruginosa* despite inducing innate immune responses, we explored how this inhibition affected gut colonization by *P. aeruginosa*. Knockdown of *eft-2* and *eef-1B.2* significantly decreased gut colonization with *P. aeruginosa* in N2 animals (Fig. 8A-C). This finding was paradoxical, as the knockdown of *eft-2* and *eef-1B.2* reduced the survival of N2 animals on *P. aeruginosa*, even though gut colonization by the pathogen was concurrently decreased. These results again suggested that gut colonization levels by *P. aeruginosa* might not be a reliable predictor of *C. elegans* survival during infection.

**Figure 8.**
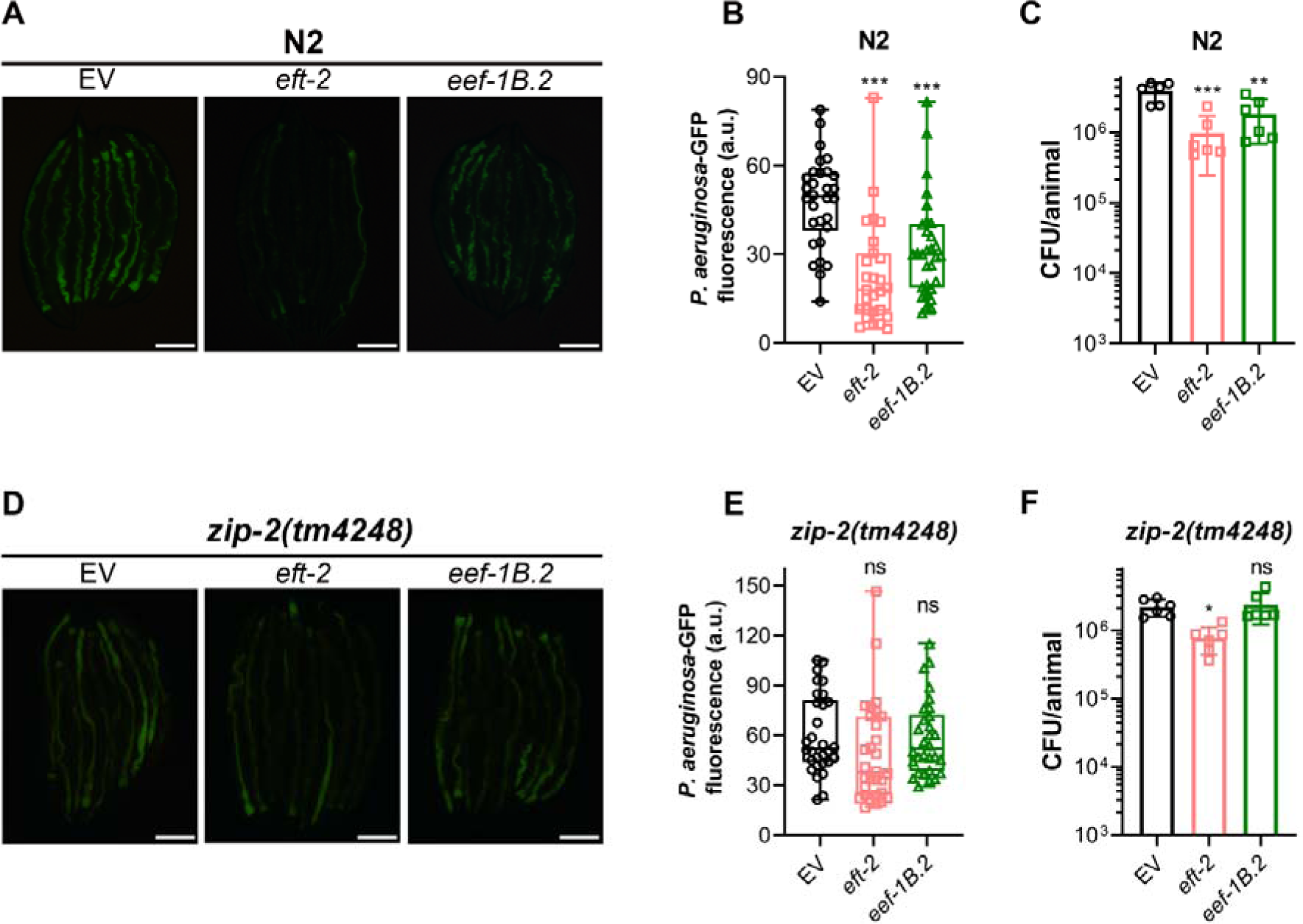
Inhibition of translation elongation reduces *C. elegans* gut colonization with *P. aeruginosa* partly via ZIP-2. (A) Representative fluorescence images of N2 animals incubated on *P. aeruginosa*-GFP for 24 hours at 25°C after treatment with the empty vector (EV) control, *eft-2*, and *eef-1B.2* RNAi. Scale bar = 200 μm. (B) Quantification of GFP levels of N2 animals incubated on *P. aeruginosa*-GFP for 24 hours at 25°C after treatment with the EV control, *eft-2*, and *eef-1B.2* RNAi. \*\*\**P* < 0.001 via ordinary one-way ANOVA followed by Dunnett’s multiple comparisons test (*n* = 30 worms for EV control and *eef-1B.2* RNAi, and 27 worms for *eft-2* RNAi). (C) Colony-forming units (CFU) per animal of N2 worms incubated on *P. aeruginosa*-GFP for 24 hours at 25°C after treatment with the EV control, *eft-2*, and *eef-1B.2* RNAi. \*\*\**P* < 0.001 and \*\**P* < 0.01 via ordinary one-way ANOVA followed by Dunnett’s multiple comparisons test (*n = 6* biological replicates). (D) Representative fluorescence images of *zip-2(tm4248)* animals incubated on *P. aeruginosa*-GFP for 24 hours at 25°C after treatment with the EV control, *eft-2*, and *eef-1B.2* RNAi. Scale bar = 200 μm. (E) Quantification of GFP levels of *zip-2(tm4248)* animals incubated on *P. aeruginosa*-GFP for 24 hours at 25°C after treatment with the EV control, *eft-2*, and *eef-1B.2* RNAi. ns, nonsignificant via ordinary one-way ANOVA followed by Dunnett’s multiple comparisons test (*n* = 30 worms each). (F) CFU per animal of *zip-2(tm4248)* worms incubated on *P. aeruginosa*-GFP for 24 hours at 25°C after treatment with the EV control, *eft-2*, and *eef-1B.2* RNAi. \**P* < 0.05 and ns, nonsignificant via ordinary one-way ANOVA followed by Dunnett’s multiple comparisons test (*n = 6* biological replicates).

Next, we evaluated whether any of the immunity pathways mediated the reduced colonization of the gut with *P. aeruginosa* upon *eft-2* and *eef-1B.2* knockdown. Inhibition of *eft-2* and *eef-1B.2* resulted in diminished *P. aeruginosa* colonization in *sek-1(km4)*, *pmk-1(km25)*, *dbl-1(nk3)*, and *nhr-8(ok186)* animals (Fig. S9). However, *eft-2* and *eef-1B.2* knockdown did not lead to a significant reduction in *P. aeruginosa* colonization in *zip-2(tm4248)* animals (Fig. 8D and E). Although there was a decent correlation between the *P. aeruginosa*-GFP levels in the gut and CFU per animal across different *C. elegans* strains (Fig. S10), a slight discrepancy was observed in *zip-2(tm4248)* animals. While *eft-2* knockdown did not significantly reduce *P. aeruginosa*-GFP levels, it did result in a small but significant reduction in CFU levels in *zip-2(tm4248)* animals (Fig. 8E and F). These findings suggested that the reduction in pathogen colonization due to translation elongation inhibition occurs largely in a ZIP-2-dependent manner. Overall, these results indicated that inhibiting translation elongation reduces *C. elegans* survival on *P. aeruginosa*, despite a decrease in gut colonization by the bacterium.

### Inhibition of translation initiation and elongation leads to overlapping and unique transcriptional changes

Finally, we examined the overlap of genes upregulated in N2 animals following *eif-2*α and *eft-2* knockdown. While 1008 genes were upregulated by both *eif-2*α and *eft-2* knockdown, 789 genes were uniquely upregulated by *eif-2*α knockdown, and 394 genes were uniquely upregulated by *eft-2* knockdown (Fig. 9A). GO analysis of genes uniquely upregulated by *eif-2*α knockdown revealed enrichment for structural constituents of the cuticle, but not for innate immune responses (Fig. 9B and C). Conversely, GO analysis of genes upregulated by both *eif-2*α and *eft-2* knockdown showed enrichment for innate immune responses (Fig. 9D and E). Interestingly, genes uniquely upregulated by *eft-2* knockdown were enriched for innate immune responses and kinase activity (Fig. 9F and G). These data showed that inhibiting translation elongation activates immune genes that are not triggered by translation initiation inhibition.

**Figure 9.**
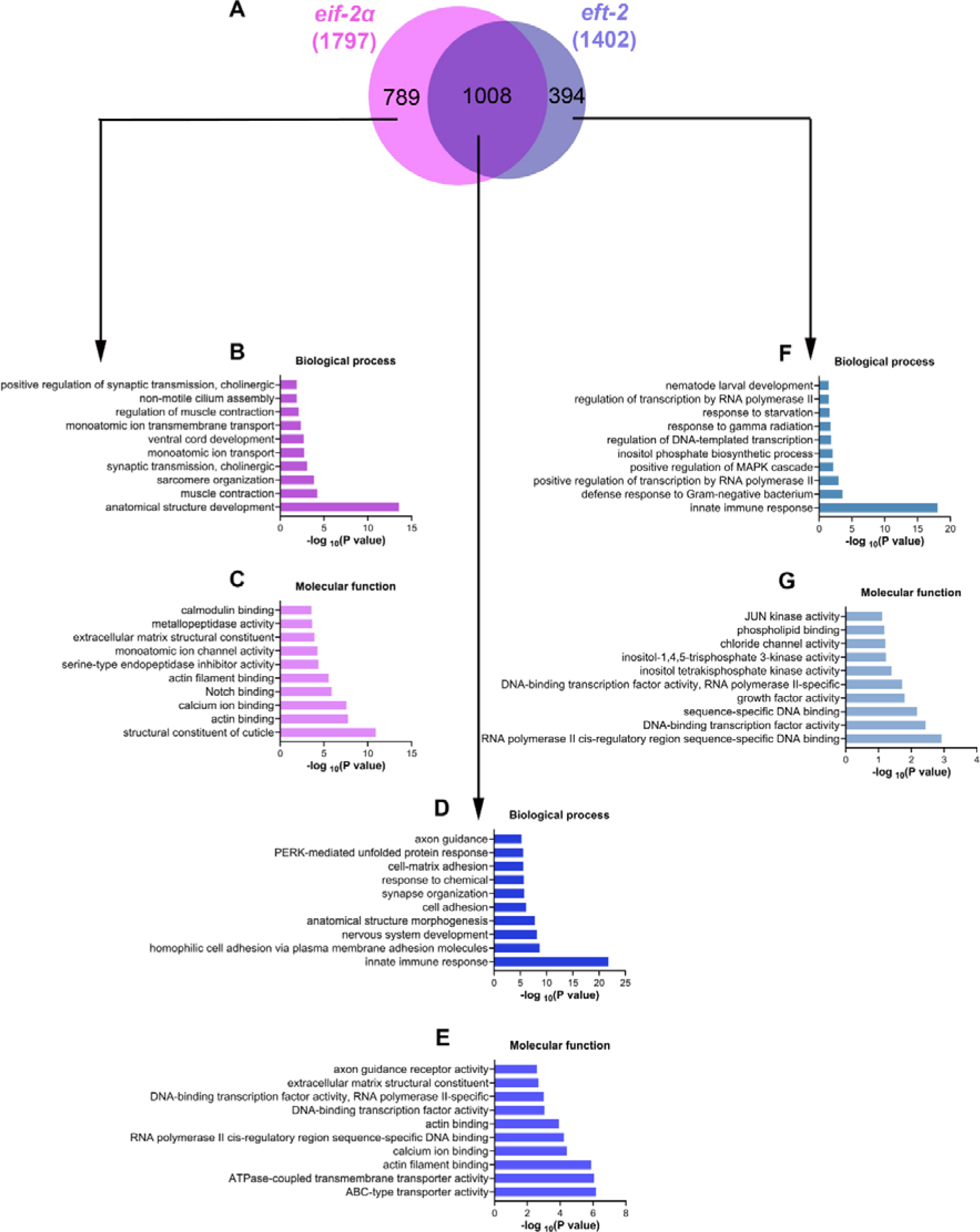
Inhibition of translation initiation and elongation results in overlapping and unique transcriptional changes. (A) Venn diagram showing the overlap between genes upregulated upon *eif-2*α and *eft-2* knockdown in N2 animals. (B)-(C) Gene Ontology (GO) enrichment analysis of genes upregulated uniquely in *eif-2*α knockdown in N2 animals for biological processes (B) and molecular functions (C). (D)-(E) GO enrichment analysis of genes upregulated both in *eif-2*α and *eft-2* knockdown in N2 animals for biological processes (D) and molecular functions (E). (F)-(G) GO enrichment analysis of genes upregulated uniquely in *eft-2* knockdown in N2 animals for biological processes (F) and molecular functions (G).

## Discussions

A large number of pathogens, including viruses and bacteria, block host translation (Mccormick and Khaperskyy, 2017; Remick et al., 2023). The dampening of host translation may assist the pathogen in evading host immune responses, given that the synthesis of host immune effector polypeptides is concurrently suppressed. Since various pathogens target host protein synthesis, the blockade of translation might simulate a pathogen attack, consequently triggering the activation of immune responses. Indeed, the inhibition of protein synthesis, independent of pathogens, has been demonstrated to induce immune responses (Dunbar et al., 2012; Govindan et al., 2015; Melo and Ruvkun, 2012). However, the mere elevation in the mRNA levels of immune genes may not serve as a reliable indicator of the host’s ability to survive the pathogen attack. For example, the *Pseudomonas entomophila*-mediated blockade of *Drosophila* protein synthesis resulted in an impaired immune response, notwithstanding the upregulation of mRNA levels of immune genes (Chakrabarti et al., 2012). Hence, it becomes imperative to couple the analysis of immune gene mRNA levels with the host’s survival upon pathogen challenge following the inhibition of translation. In the current study, we demonstrate that, despite the increased mRNA levels of immune genes, inhibition of translation initiation and elongation can yield opposing outcomes in terms of host survival when challenged with a pathogen.

We have shown that inhibiting translation initiation enhances survival, while inhibiting translation elongation diminishes the survival of *C. elegans* on *P. aeruginosa*. Why would inhibition of translation initiation and elongation yield contrasting effects on *C. elegans* survival on *P. aeruginosa* when both interventions lead to increased mRNA levels of immune genes? One plausible explanation is that the inhibition of the elongation step produces toxic premature polypeptides (Svidritskiy et al., 2018), while inhibition of the initiation step decreases overall protein synthesis without generating premature peptides. However, since the adverse effects of inhibiting the elongation step are fully rescued by the knockdown of ZIP-2, it seems unlikely that reduced survival results from toxic byproducts. If toxic premature polypeptides were the primary cause, blocking ZIP-2-mediated immune responses would not be expected to reverse these adverse effects. Another possibility is that the survival on the pathogen hinges on the balance between increased energy consumption due to elevated mRNA synthesis for immune genes and the quantity of immune effector polypeptides produced upon inhibiting translation initiation and elongation. Nevertheless, since inhibiting both initiation and elongation factors reduces *C. elegans* gut colonization with *P. aeruginosa*, an effective immune response is likely generated when both steps are inhibited. In future studies, it will be useful to carry out proteomics analysis upon inhibition of translation initiation and elongation and compare it with the transcriptome changes to understand if specific immune-related polypeptides are upregulated or downregulated upon inhibition of translation initiation and elongation.

It is also conceivable that inhibiting translation elongation triggers a hyperimmune response, which proves detrimental to the host despite reducing gut colonization with *P. aeruginosa*. Indeed, inhibition of translation elongation has been shown to induce more robust immune responses than inhibition of initiation (Dunbar et al., 2012). Our RNA sequencing data demonstrated that inhibiting translation elongation activates immune genes that are not activated by the inhibition of translation initiation. The activation of hyperimmune responses has been shown to reduce the survival of *C. elegans* on *P. aeruginosa* despite diminishing gut colonization by the bacterium (Rao et al., 2024). Given that the knockout of ZIP-2 reverses the adverse effects of inhibiting translation elongation on host survival, a hyperimmune response could underlie the detrimental consequences of inhibiting translation elongation. Because translation elongation inhibition activates ZIP-2-dependent and ZIP-2-independent immune responses, knockout of ZIP-2 might dampen the hyperimmune response triggered by translation elongation inhibition. Notably, while the knockdown of *eft-2* has detrimental effects in wild-type N2 animals, it proves beneficial in several innate immunity mutants. Thus, the aberrant activation of immune responses upon translation elongation inhibition may benefit immunocompromised animals. Given that translation elongation inhibition activates ZIP-2-independent immune responses and that ZIP-2-independent genes are enriched in protein kinase activity, it is plausible that ZIP-2-independent immune responses are mediated by protein kinases. Indeed, a previous study demonstrated that multiple kinases regulate surveillance immunity upon translation elongation inhibition (Govindan et al., 2015). Future research is needed to elucidate the additional mechanisms underlying the reduced survival of *C. elegans* on *P. aeruginosa* when translation elongation is inhibited.

A noteworthy difference between the genes upregulated by translation initiation and elongation inhibition involves the collagen genes. Inhibition of translation initiation resulted in a ZIP-2-dependent enrichment of collagen genes, which are constituents of the cuticle. Interestingly, genes upregulated by translation elongation inhibition were not enriched in collagens. Increased expression of collagens can improve *C. elegans* lifespan (Ewald et al., 2015) and survival on *P. aeruginosa* (Sellegounder et al., 2019). Therefore, it is possible that the increased expression of collagens contributes to the improved survival of animals on *P. aeruginosa* upon translation initiation inhibition.

Innate immunity appears to be a pivotal factor for enhanced lifespan (Garsin et al., 2003; Soo et al., 2023). Because the inhibition of translation contributes to an extended lifespan, it becomes crucial to discern whether the increased lifespan resulting from translation blockage is attributable to an augmented immune response. Inhibition of both translation initiation and elongation is known to increase *C. elegans* lifespan (Li et al., 2011; Statzer et al., 2022). However, our findings reveal a dichotomy: while the inhibition of translation initiation enhances survival, the inhibition of translation elongation reduces the survival of *C. elegans* on *P. aeruginosa*. Notably, the knockdown of *hel-1*, functioning as both an RNA helicase and a translation initiation factor, is known to reduce *C. elegans* lifespan by about 9% (Seo et al., 2015). On the other hand, our observations indicate that its knockdown extends *C. elegans* survival on *P. aeruginosa* by approximately 33% (Fig. 1G). The translation initiation factor *ife-2*, known for increasing lifespan by about 18% (Matei et al., 2021), exhibits a distinct pattern. Although it extends lifespan, it does not alter the survival of *C. elegans* on *P. aeruginosa* (Fig. 1G). These studies imply that pathways related to immunity may not necessarily mediate the increased lifespan resulting from translation inhibition. While ZIP-2 appears to be the major immune modulator when translation initiation is inhibited, several pathways work parallelly during translation initiation inhibition to modulate lifespan (Hansen et al., 2007b; Howard et al., 2016; Li et al., 2011; Pan et al., 2007; Rogers et al., 2011). These observations suggest that translation inhibition-mediated longevity and pathogen resistance are regulated by distinct genetic mechanisms.

## Materials and methods

### Bacterial strains

The bacterial strains utilized in this study include *Escherichia coli* OP50, *E. coli* HT115(DE3), *Pseudomonas aeruginosa* PA14, *P. aeruginosa* PA14 expressing green fluorescent protein (*P. aeruginosa* PA14-GFP). The GFP expression in *P. aeruginosa* PA14-GFP is from plasmid pRR54GFP19-1, which is described earlier (Tan et al., 1999). The cultures of *E. coli* OP50, *E. coli* HT115(DE3), and *P. aeruginosa* PA14 were grown in Luria-Bertani (LB) broth at 37°C, while the cultures of *P. aeruginosa* PA14-GFP were grown in LB broth with 50 µg/mL kanamycin at 37°C.

### *C. elegans* strains and growth conditions

*C. elegans* hermaphrodites were maintained at 20°C on nematode growth medium (NGM) plates seeded with *E. coli* OP50 as the food source unless otherwise indicated. The wild-type control used was Bristol N2. The following strains were used in the study: *zip-2(tm4248),* KU25 *pmk-1(km25),* NU3 *dbl-1(nk3),* AE501 *nhr-8(ok186),* KU4 *sek-1(km4)*, and HC196 *sid-1(qt9)*. All the strains were obtained from the Caenorhabditis Genetics Center (University of Minnesota, Minneapolis, MN) except for *zip-2(tm4248*), which was obtained from the National BioResearch Project (*C. elegans*), Japan.

### *C. elegans* development assays

Wild-type N2 gravid hermaphrodites were lysed in an alkaline bleach solution to obtain eggs, which were then washed three times with the M9 buffer solution. The eggs were then incubated in the M9 buffer solution for 22 hours at room temperature in a 1.5 mL microcentrifuge tube to obtain synchronized L1-arrested worms. These synchronized L1 worms were then transferred to RNAi plates targeting different translation factors and incubated at 20°C for 72 hours. Following this, animals were scored for normal or slowed development. At least three independent experiments (biological replicates) were performed for each condition.

### RNA interference (RNAi)

RNAi was employed to induce loss-of-function phenotypes by feeding worms with the *E. coli* strain HT115(DE3) expressing double-stranded RNA homologous to a target *C. elegans* gene. The list of *C. elegans* genes encoding translation factors was obtained from (Rhoads et al., 2006). RNAi was carried out as described previously (Das et al., 2023). Briefly, *E. coli* HT115(DE3) with the appropriate vectors were grown in LB broth containing ampicillin (100 μg/mL) at 37°C overnight on a shaker, concentrated ten times, and plated onto RNAi NGM plates containing 100 μg/mL ampicillin and 3 mM isopropyl β-D-thiogalactoside (IPTG). The plated bacteria were allowed to grow overnight at 37°C. Out of 47 translation factors, the knockdown of 23 led to the slowed development of worms. Therefore, for the genes whose knockdown led to slowed development (Table S1), the worms were grown on the empty vector control RNAi until the L3 stage before being transferred to the corresponding gene RNAi. The worms were grown on RNAi bacteria till the 1-day-adult stage before being utilized for subsequent experiments. The eggs were allowed to develop at 20°C for 96 hours on RNAi for the genes whose knockdown did not affect development, except for *eef-1B.2*. For comparison with *eft-2*, RNAi against *eef-1B.*2 was initiated from the L3 stage. The RNAi clones were from the Ahringer RNAi library and were verified by sequencing.

### C. elegans killing assays on P. aeruginosa PA14

The full-lawn killing assays of *C. elegans* on *P. aeruginosa* PA14 were conducted as described earlier (Singh and Aballay, 2019a). Briefly, bacterial cultures were prepared by inoculating individual bacterial colonies of *P. aeruginosa* into 3 mL of LB and grown for 10-12 hours on a shaker at 37°C. Bacterial lawns were prepared by spreading 20 µL of the culture on the entire surface of 3.5-cm-diameter modified NGM agar plates (0.35% instead of 0.25% peptone). The plates were incubated at 37°C for 10-12 hours and then cooled to room temperature for at least 30 minutes before seeding with synchronized 1-day-old adult animals. The killing assays were carried out at 25°C, and live animals were transferred daily to fresh *P. aeruginosa* plates. Animals were scored at times indicated and were considered dead when they failed to respond to touch. At least three independent experiments (biological replicates) were performed for each condition.

### *P. aeruginosa*-GFP colonization assay

The *P. aeruginosa* PA14-GFP colonization assays were carried out as described earlier (Rao et al., 2024; Singh and Aballay, 2019b). Briefly, bacterial cultures were prepared by inoculating individual bacterial colonies of *P. aeruginosa* PA14-GFP into 3 mL of LB containing 50 μg/mL kanamycin and grown for 10-12 hours on a shaker at 37°C. Bacterial lawns were prepared by spreading 20 µL of the culture on the entire surface of 3.5-cm-diameter modified NGM agar plates (0.35% instead of 0.25% peptone) containing 50 μg/mL of kanamycin. The plates were incubated at 37°C for 12 hours and then cooled to room temperature for at least 30 minutes before seeding with 1-day-old adult gravid adults. The assays were performed at 25°C. At indicated times, the worms were picked under a non-fluorescence stereomicroscope and visualized within 5 minutes under a fluorescence microscope. At least three independent experiments (biological replicates) were performed for each condition.

### Quantification of intestinal bacterial loads

The intestinal *P. aeruginosa* PA14-GFP bacterial loads were quantified by measuring colony-forming units (CFU) as described earlier (Das et al., 2023; Rao et al., 2024). Briefly, lawns of *P. aeruginosa* PA14-GFP were prepared as described above. At the indicated times for each experiment, the animals were transferred from *P. aeruginosa*-GFP plates to fresh *E. coli* OP50 plates for approximately 10 minutes to eliminate bacteria stuck to their body. The animals were transferred to fresh *E. coli* OP50 plates two times more, with each transfer lasting approximately 10 minutes. Subsequently, ten animals per condition were transferred into 50 μL of PBS containing 0.01% triton X-100 and lysed using glass beads. Serial dilutions of the lysates (10^1^, 10^2^, 10^3^, 10^4^, and 10^5^) were seeded on LB plates containing 50 μg/mL of kanamycin to select for *P. aeruginosa*-GFP cells and grown overnight at 37°C. Single colonies were counted the next day and represented as the number of bacterial cells or CFU per animal. Six independent experiments were performed for each condition.

### RNAi efficiency test for *zip-2(tm4248)*

The wild-type N2 and *zip-2(tm4248)* animals were synchronized by egg laying on *act-5* and *bli-3* RNAi plates along with the empty vector control RNAi plates. The *sid-1(qt9)* animals were used as RNAi-defective controls (Whangbo et al., 2017). The plates with eggs were incubated at 20°C for 72 hours. After that, the animals were monitored for development defects and blisters on the cuticle for *act-5* and *bli-3* RNAi, respectively.

### *C. elegans* RNA isolation and quantitative reverse transcription-PCR (qRT-PCR)

Both wild-type N2 and *zip-2(tm4248)* animals were synchronized by egg-laying. Approximately 50 gravid adult animals were transferred to 9-cm control empty vector RNAi plates and allowed to lay eggs for 4-hours. The gravid adults were then removed, and the eggs were allowed to develop at 20°C for 36 hours (until the L3 stage). After that, the synchronized L3 animals were collected in M9, washed twice with M9, and transferred to 9-cm *eif-2*α*, ifg-1, eft-2*, and *eef-1B.2* RNAi plates and were allowed to grow till 1-day-old adults. The control animals were maintained on the empty vector RNAi till they grew to 1-day-old adults. Subsequently, the animals were collected, washed with M9 buffer, and frozen in TRIzol reagent (Life Technologies, Carlsbad, CA). Total RNA was extracted using the RNeasy Plus Universal Kit (Qiagen, Netherlands). A total of 1 μg of total RNA was reverse-transcribed with random primers using the PrimeScript™ 1st strand cDNA Synthesis Kit (TaKaRa) according to the manufacturer’s protocols. qRT-PCR was conducted using TB Green fluorescence (TaKaRa) on a MasterCycler EP Realplex 4 thermal cycler (Eppendorf) in 96-well-plate format. Fifteen microliter reactions were analyzed as outlined by the manufacturer (TaKaRa). The relative fold-changes of the transcripts were calculated using the comparative *CT*(2^-ΔΔ^*CT*) method and normalized to pan-actin (*act-1, -3, -4*) as described earlier (Singh and Aballay, 2017). All samples were run in triplicate (technical replicates) and repeated at least four times (biological replicates). The sequence of the primers is provided in Table S10.

### RNA sequencing and data analysis

Wild-type N2 and *zip-2(tm4248)* animals were exposed to *eif-2*α and *eft-*2 RNAi treatments, along with empty vector control RNAi, as described above. Total RNA samples were isolated from three biological replicates, as described above. Library preparation and sequencing were conducted at Unipath Specialty Laboratory Ltd, India. The cDNA libraries were sequenced on the NovaSeq 6000 platform, using 150 bp paired-end reads.

The RNA sequencing data were analyzed using the web platform Galaxy (https://usegalaxy.org/) as described earlier (Rao et al., 2024). Briefly, the paired-end reads were first trimmed using the Trimmomatic tool. The trimmed reads were then mapped to the *C. elegans* genome (WS220) using the aligner STAR. The number of reads mapped to each gene was counted using the *htseq-count* tool. Differential gene expression analysis was then performed using DESeq2. Genes exhibiting at least two-fold change and *P*-value <0.01 were considered differentially expressed. Gene Ontology analysis was performed using the DAVID Bioinformatics Database (https://david.ncifcrf.gov/tools.jsp). Venn diagrams were generated using the web tool BioVenn (https://www.biovenn.nl/) (Hulsen et al., 2008).

### Fluorescence imaging of *C. elegans*

Fluorescence imaging was carried out as described previously (Gokul and Singh, 2022; Ravi et al., 2023). Briefly, the animals were picked under a non-fluorescence stereomicroscope to avoid potential bias. The animals were anesthetized using an M9 salt solution containing 50 mM sodium azide and mounted onto 2% agarose pads. The animals were then visualized using a Nikon SMZ-1000 fluorescence stereomicroscope. The fluorescence intensity was quantified using Image J software.

### Quantification and statistical analysis

The statistical analysis was performed with Prism 8 (GraphPad). All error bars represent the mean ± standard deviation (SD). The unpaired, two-tailed, two-sample *t*-test was employed when needed, and statistical significance was determined when *P* < 0.05. For more than two samples, ordinary one-way ANOVA followed by Dunnett’s multiple comparisons test was used. In the figures, asterisks (*) denote statistical significance as follows: *, *P* < 0.05, **, *P* < 0.01, ***, *P* < 0.001, as compared with the appropriate controls. The Kaplan-Meier method was utilized to calculate the survival fractions, and statistical significance between survival curves was determined using the log-rank test. All experiments were performed at least three times unless otherwise indicated.

## Acknowledgments

We thank the Caenorhabditis Genetics Center (funded by the NIH Office of Research Infrastructure Programs (P40 OD010440)) and the National BioResearch Project (*C. elegans*), Japan, for providing the strains used in this study.

## Data availability

The RNA sequencing data for N2 and *zip-2(tm4248)* worms grown each on empty vector control, *eif-2*α, and *eft-2* RNAi have been submitted to the public repository, the Sequence Read Archive, with BioProject ID PRJNA1145895. All data generated or analyzed during this study are included in the manuscript and supporting files.

## Supplementary Figures

**Figure S1.**
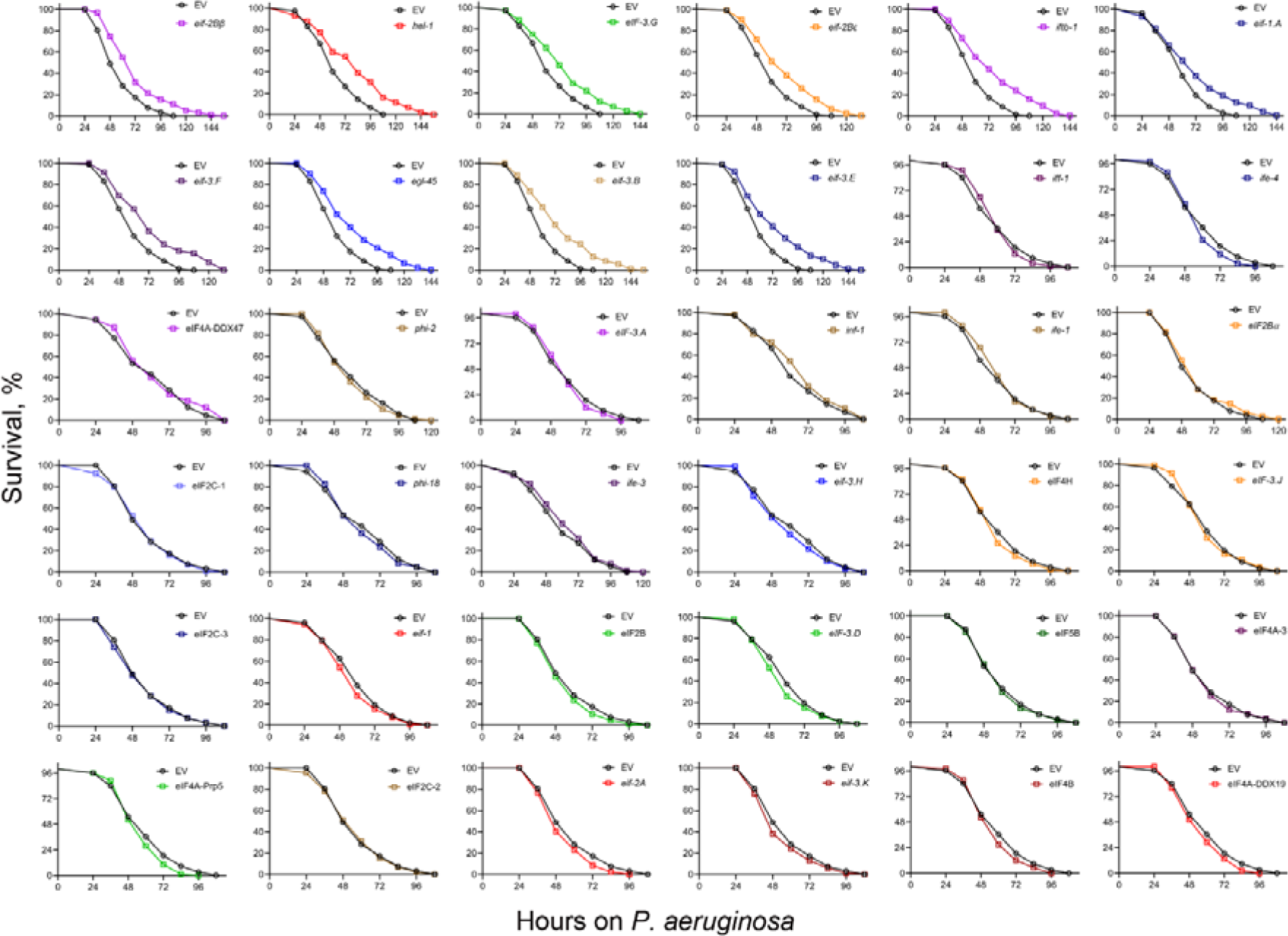
Inhibition of several translation initiation factors improves *C. elegans* survival on *P. aeruginosa*. Representative survival plots of N2 animals on *P. aeruginosa* PA14 at 25°C after treatment with the empty vector (EV) control and RNAi against various translation initiation factors. The EV control is common for different cohorts of the survival curves. Detailed statistical analysis is available in Table S2.

**Figure S2.**
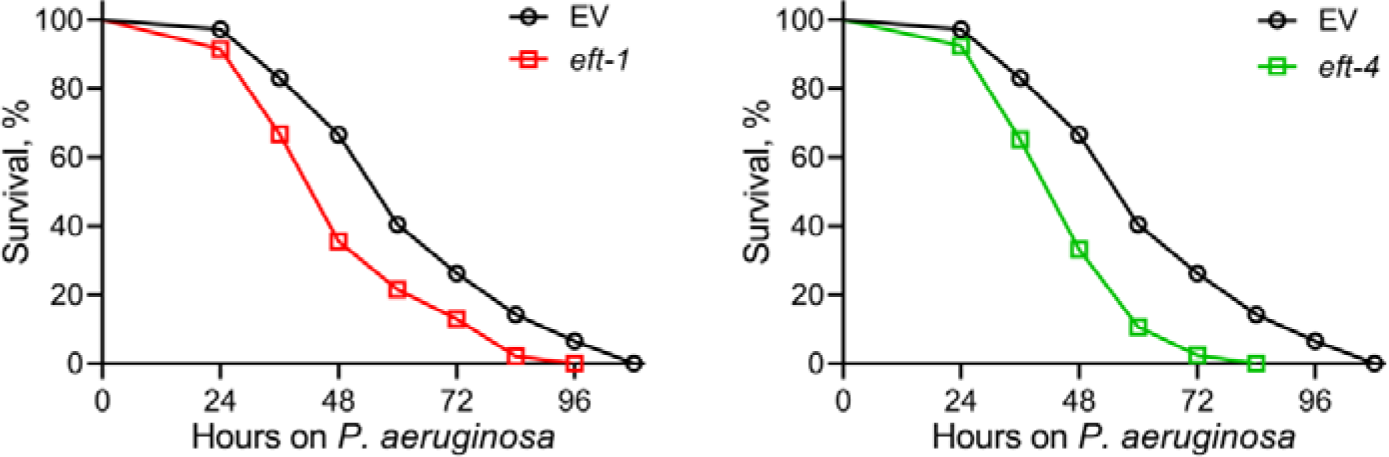
Inhibition of translation elongation factors reduces *C. elegans* survival on *P. aeruginosa*. Representative survival plots of N2 animals on *P. aeruginosa* PA14 at 25°C after treatment with the empty vector (EV) control, *eft-1*, and *eft-4* RNAi. The EV control is common for the two survival plots. Detailed statistical analysis is available in Table S2.

**Figure S3.**
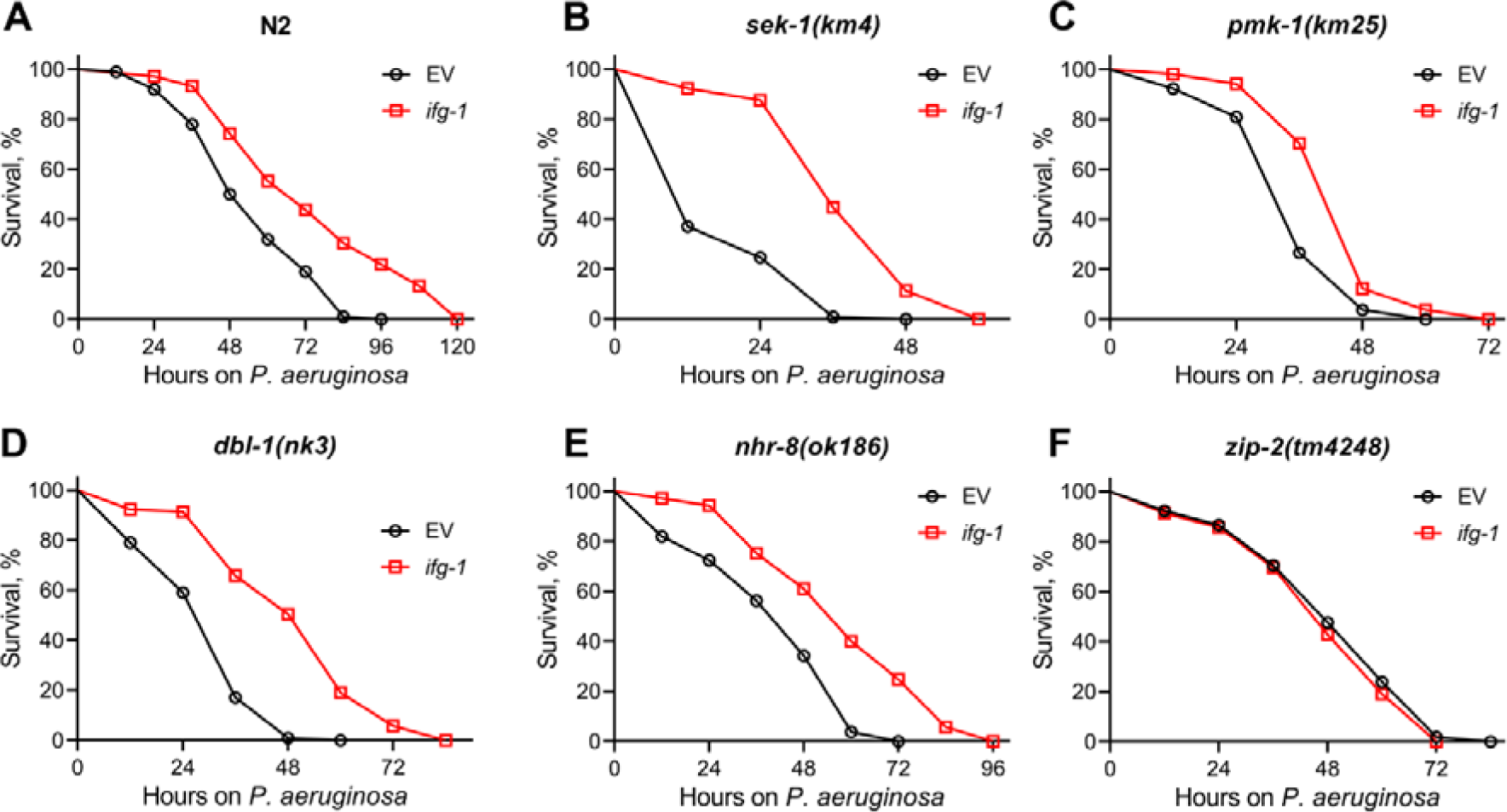
Knockdown of *ifg-1* increases *C. elegans* survival on *P. aeruginosa* via transcription factor ZIP-2. Representative survival plots of N2 (A), *sek-1(km4)* (B), *pmk-1(km25)* (C), *dbl-1(nk3)* (D), *nhr-8(ok186)* (E), and *zip-2(tm4248)* (F) animals on *P. aeruginosa* PA14 at 25°C after treatment with the empty vector (EV) control and *ifg-1* RNAi. *P*<0.001 for *ifg-1* RNAi compared to EV control for N2, *sek-1(km4)*, *pmk-1(km25)*, *dbl-1(nk3)*, and *nhr-8(ok186)* animals. Survival curves for *ifg-1* RNAi compared to the EV control for *zip-2(tm4248)* animals are nonsignificant.

**Figure S4.**
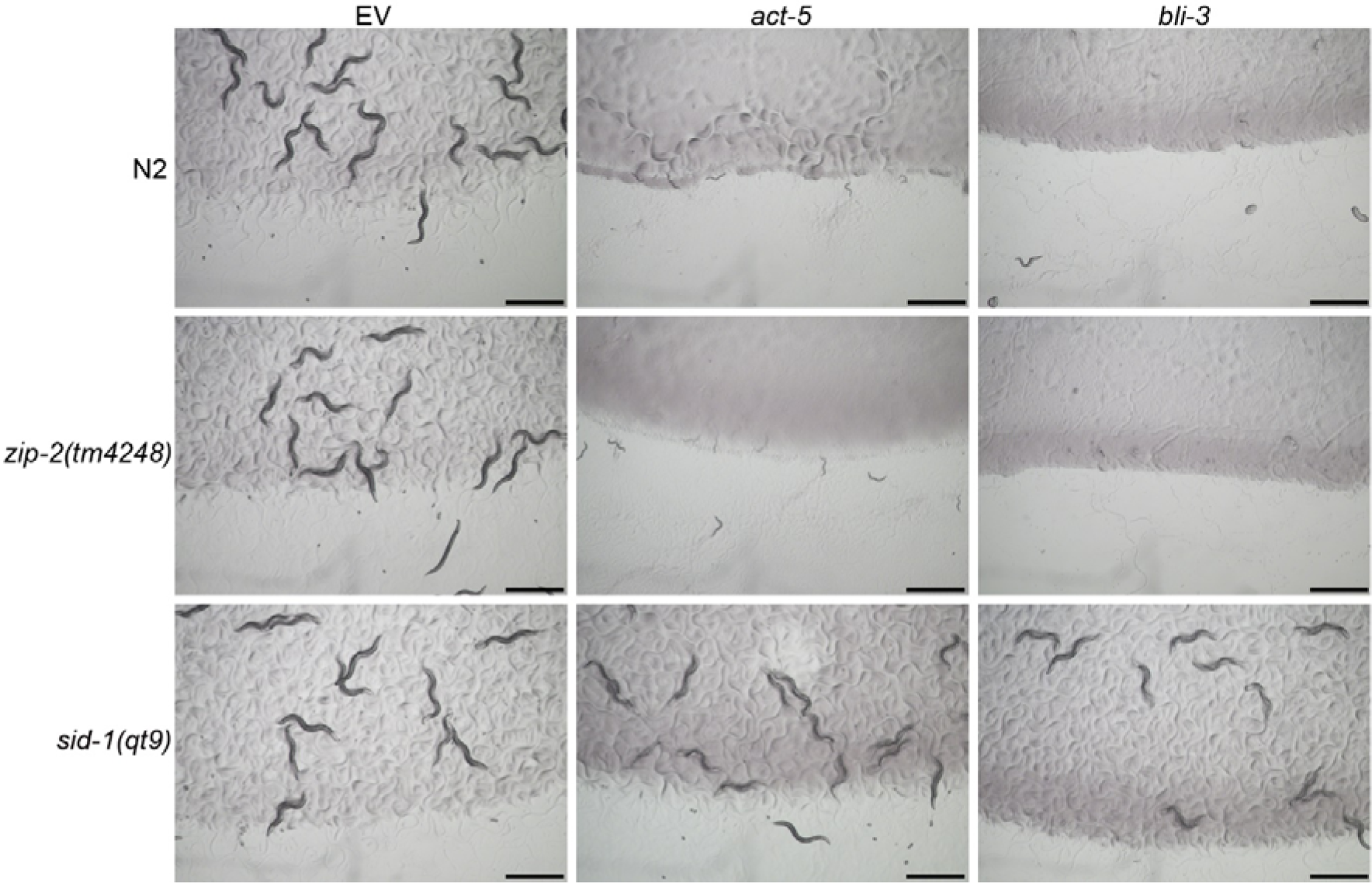
The *zip-2(tm4248)* animals are sensitive to RNAi. Representative images of wild-type N2, *zip-2(tm4248)*, and *sid-1(qt9)* worms on the empty vector (EV) control, *act-5*, and *bli-3* RNAi. Scale bar = 1 mm.

**Figure S5.**
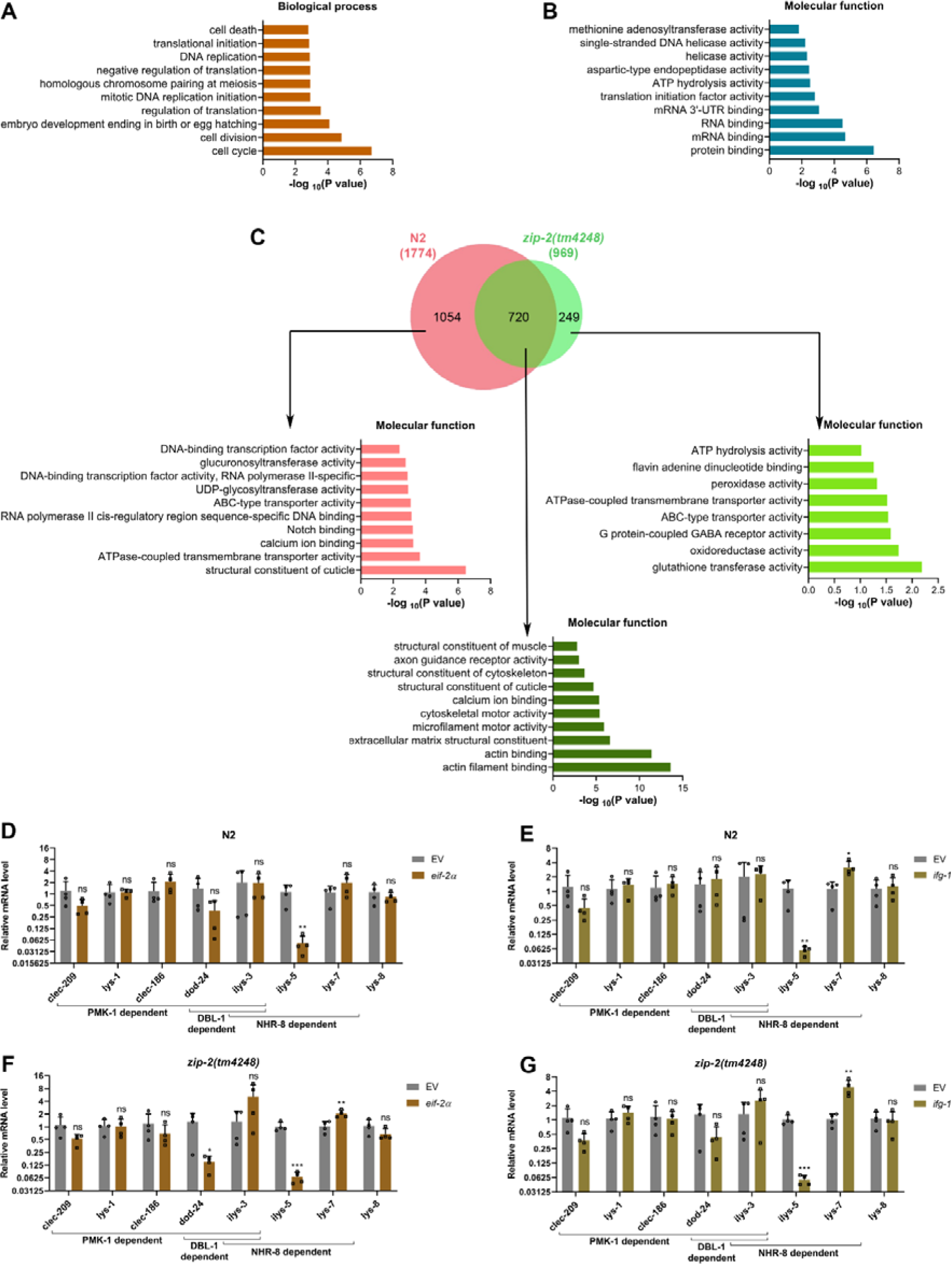
Inhibition of translation initiation activates a ZIP-2-dependent immune response. (A)-(B) Gene Ontology (GO) enrichment analysis of downregulated genes upon *eif-2*α knockdown in N2 animals for biological processes (A) and molecular functions (B). (C) Venn diagram showing the overlap between genes upregulated upon *eif-2*α knockdown in N2 and *zip-2(tm4248)* animals. The GO analysis for molecular functions of unique and common genes is shown. (D) Quantitative reverse transcription-PCR (qRT-PCR) for immune genes expression analysis of N2 animals after treatment with the empty vector (EV) control and *eif-2*α RNAi. (E) qRT-PCR for immune genes expression analysis of N2 animals after treatment with the EV control and *ifg-1* RNAi. (F) qRT-PCR for immune genes expression analysis of *zip-2(tm4248)* animals after treatment with the EV control and *eif-2*α RNAi. (G) qRT-PCR for immune genes expression analysis of *zip-2(tm4248)* animals after treatment with the EV control and *ifg-1* RNAi. For panels (D)-(G), \*\*\**P* < 0.001, \*\**P* < 0.01, and \**P* < 0.05 via the *t* test. ns, nonsignificant. Data represent the mean and standard deviation from four independent experiments.

**Figure S6.**
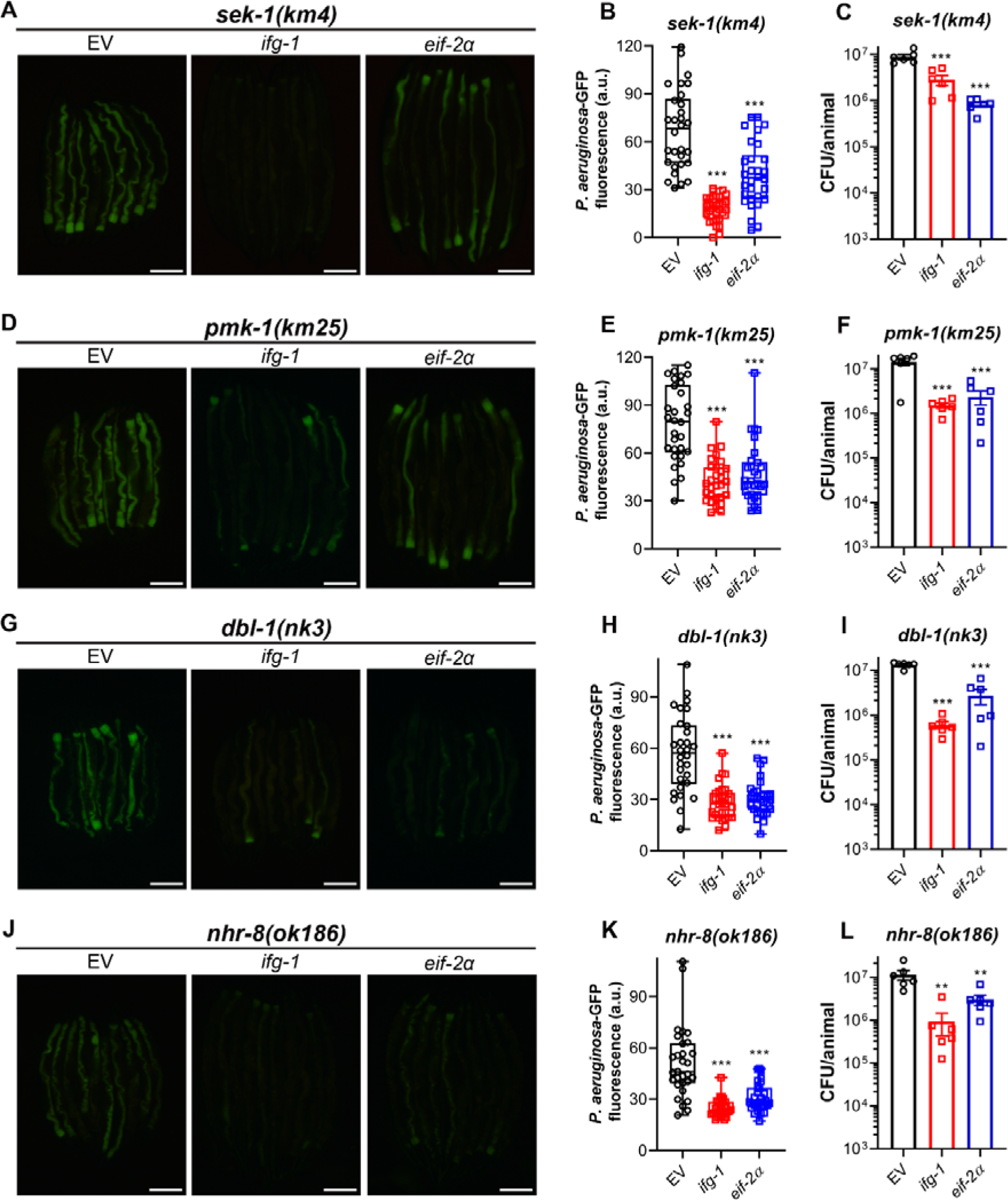
Inhibition of translation initiation reduces *P. aeruginosa* colonization in *sek-1(km4)*, *pmk-1(km25)*, *dbl-1(nk3)*, and *nhr-8(ok186)* animals. (A) Representative fluorescence images of *sek-1(km4)* animals incubated on *P. aeruginosa*-GFP for 12 hours at 25°C after treatment with the empty vector (EV) control, *ifg-1*, and *eif-2*α RNAi. Scale bar = 200 μm. (B) Quantification of GFP levels of *sek-1(km4)* animals incubated on *P. aeruginosa*-GFP for 12 hours at 25°C after treatment with the EV control, *ifg-1*, and *eif-2*α RNAi. \*\*\**P* < 0.001 via ordinary one-way ANOVA followed by Dunnett’s multiple comparisons test (*n* = 30 worms each). (C) Colony-forming units (CFU) per animal of *sek-1(km4)* animals incubated on *P. aeruginosa*-GFP for 12 hours at 25°C after treatment with the EV control, *ifg-1*, and *eif-2*α RNAi. \*\*\**P* < 0.001 via ordinary one-way ANOVA followed by Dunnett’s multiple comparisons test (*n = 6* biological replicates). (D) Representative fluorescence images of *pmk-1(km25)* animals incubated on *P. aeruginosa*-GFP for 24 hours at 25°C after treatment with the EV control, *ifg-1*, and *eif-2*α RNAi. Scale bar = 200 μm. (E) Quantification of GFP levels of *pmk-1(km25)* animals incubated on *P. aeruginosa*-GFP for 24 hours at 25°C after treatment with the EV control, *ifg-1*, and *eif-2*α RNAi. \*\*\**P* < 0.001 via ordinary one-way ANOVA followed by Dunnett’s multiple comparisons test (*n* = 30 worms for EV control and *eif-2*α RNAi, and 29 worms for *ifg-1* RNAi). (F) CFU per animal of *pmk-1(km25)* animals incubated on *P. aeruginosa*-GFP for 24 hours at 25°C after treatment with the EV control, *ifg-1*, and *eif-2*α RNAi. \*\*\**P* < 0.001 via ordinary one-way ANOVA followed by Dunnett’s multiple comparisons test (*n = 6* biological replicates). (G) Representative fluorescence images of *dbl-1(nk3)* animals incubated on *P. aeruginosa*-GFP for 24 hours at 25°C after treatment with the EV control, *ifg-1*, and *eif-2*α RNAi. Scale bar = 200 μm. (H) Quantification of GFP levels of *dbl-1(nk3)* animals incubated on *P. aeruginosa*-GFP for 24 hours at 25°C after treatment with the EV control, *ifg-1*, and *eif-2*α RNAi. \*\*\**P* < 0.001 via ordinary one-way ANOVA followed by Dunnett’s multiple comparisons test (*n* = 30 worms each). (I) CFU per animal of *dbl-1(nk3)* animals incubated on *P. aeruginosa*-GFP for 24 hours at 25°C after treatment with the EV control, *ifg-1*, and *eif-2*α RNAi. \*\*\**P* < 0.001 via ordinary one-way ANOVA followed by Dunnett’s multiple comparisons test (*n = 6* biological replicates). (J) Representative fluorescence images of *nhr-8(ok186)* animals incubated on *P. aeruginosa*-GFP for 24 hours at 25°C after treatment with the EV control, *ifg-1*, and *eif-2*α RNAi. Scale bar = 200 μm. (K) Quantification of GFP levels of *nhr-8(ok186)* animals incubated on *P. aeruginosa*-GFP for 24 hours at 25°C after treatment with the EV control, *ifg-1*, and *eif-2*α RNAi. \*\*\**P* < 0.001 via ordinary one-way ANOVA followed by Dunnett’s multiple comparisons test (*n* = 30 worms each). (L) CFU per animal of *nhr-8(ok186)* animals incubated on *P. aeruginosa*-GFP for 24 hours at 25°C after treatment with the EV control, *ifg-1*, and *eif-2*α RNAi. \*\**P* < 0.01 via ordinary one-way ANOVA followed by Dunnett’s multiple comparisons test (*n = 6* biological replicates).

**Figure S7.**
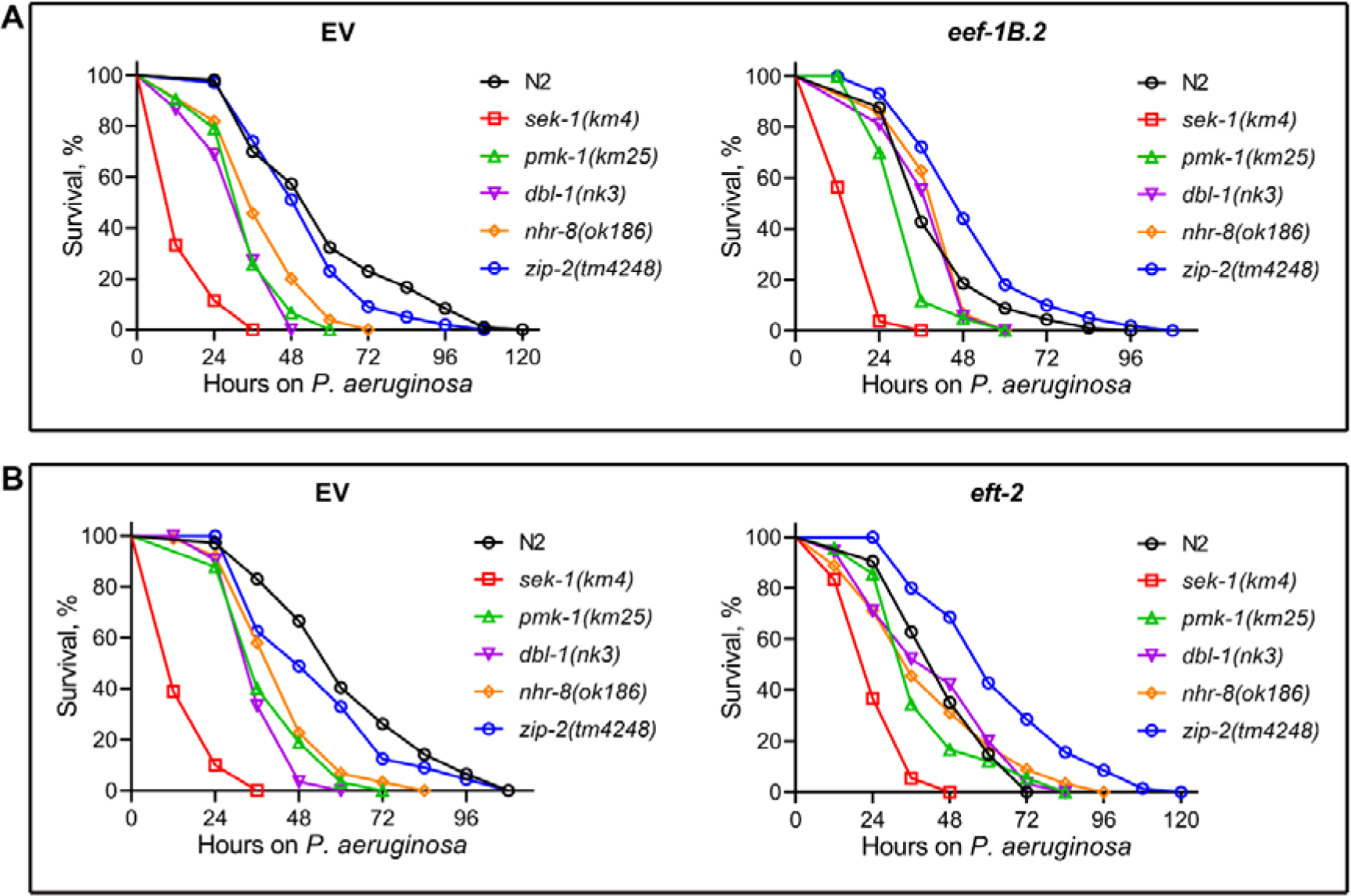
Detrimental effects of knockdown of *eef-1B.2* and *eft-2* on *C. elegans* survival on *P. aeruginosa* diminish in *zip-2(tm4248)* animals. (A) Representative survival plots of worm strains on *P. aeruginosa* PA14 at 25°C after treatment with the empty vector (EV) control RNAi (left panel) and *eef-1B.2* RNAi (right panel). (B) Representative survival plots of worm strains on *P. aeruginosa* PA14 at 25°C after treatment with the EV control RNAi (left panel) and *eft-2* RNAi (right panel).

**Figure S8.**
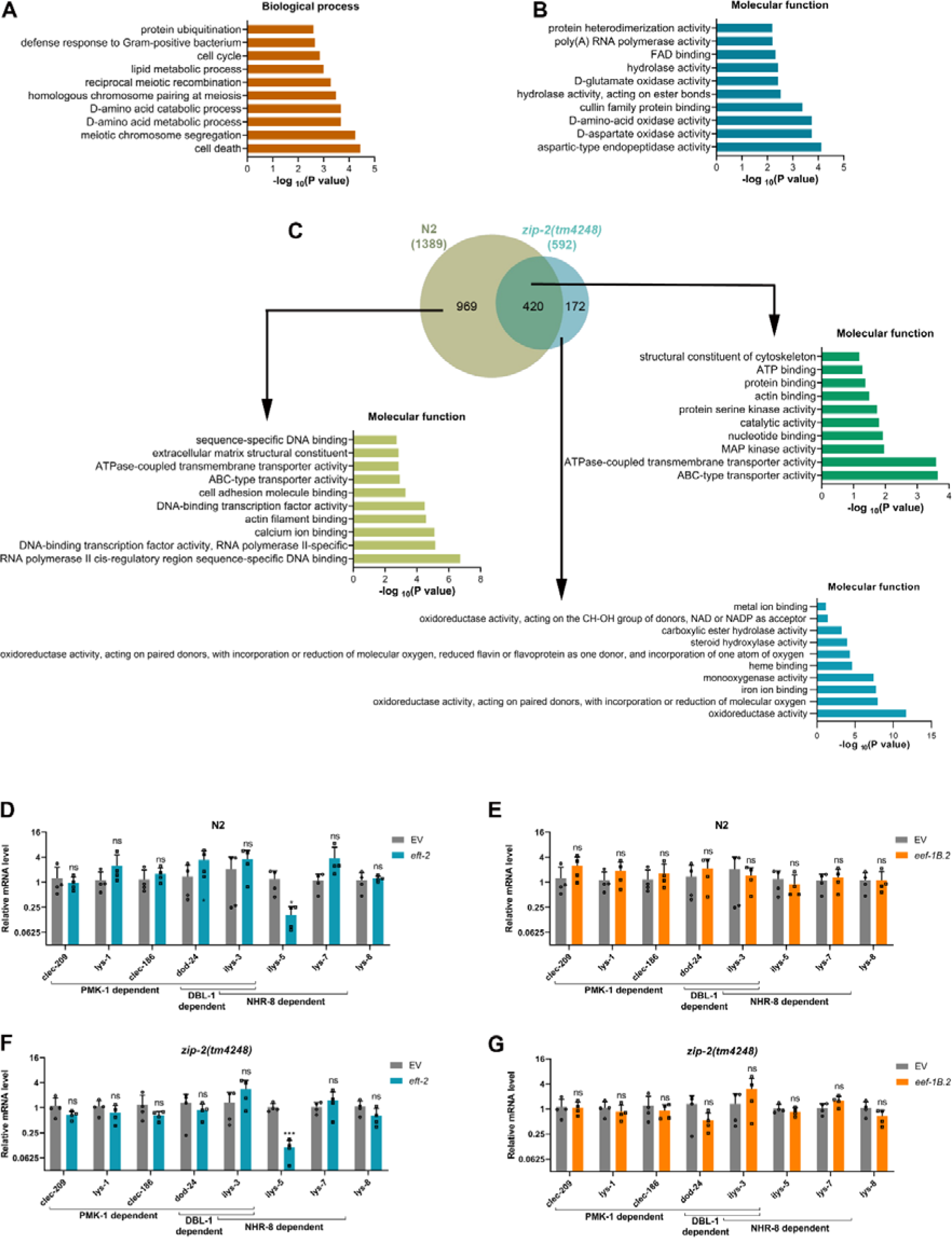
Inhibition of translation elongation activates ZIP-2-dependent and ZIP-2-independent immune responses. (A)-(B) Gene Ontology (GO) enrichment analysis of downregulated genes upon *eft-2* knockdown in N2 animals for biological processes (A) and molecular functions (B). (C) Venn diagram showing the overlap between genes upregulated upon *eft-2* knockdown in N2 and *zip-2(tm4248)* animals. The GO analysis for molecular functions of unique and common genes is shown. (D) Quantitative reverse transcription-PCR (qRT-PCR) for immune genes expression analysis of N2 animals after treatment with the empty vector (EV) control and *eft-2* RNAi. (E) qRT-PCR for immune genes expression analysis of N2 animals after treatment with the EV control and *eef-1B.2* RNAi. (F) qRT-PCR for immune genes expression analysis of *zip-2(tm4248)* animals after treatment with the EV control and *eft-2* RNAi. (G) qRT-PCR for immune genes expression analysis of *zip-2(tm4248)* animals after treatment with the EV control and *eef-1B.2* RNAi. For panels (D)-(G), \*\*\**P* < 0.001 and \**P* < 0.05 via the *t* test. ns, nonsignificant. Data represent the mean and standard deviation from four independent experiments.

**Figure S9.**
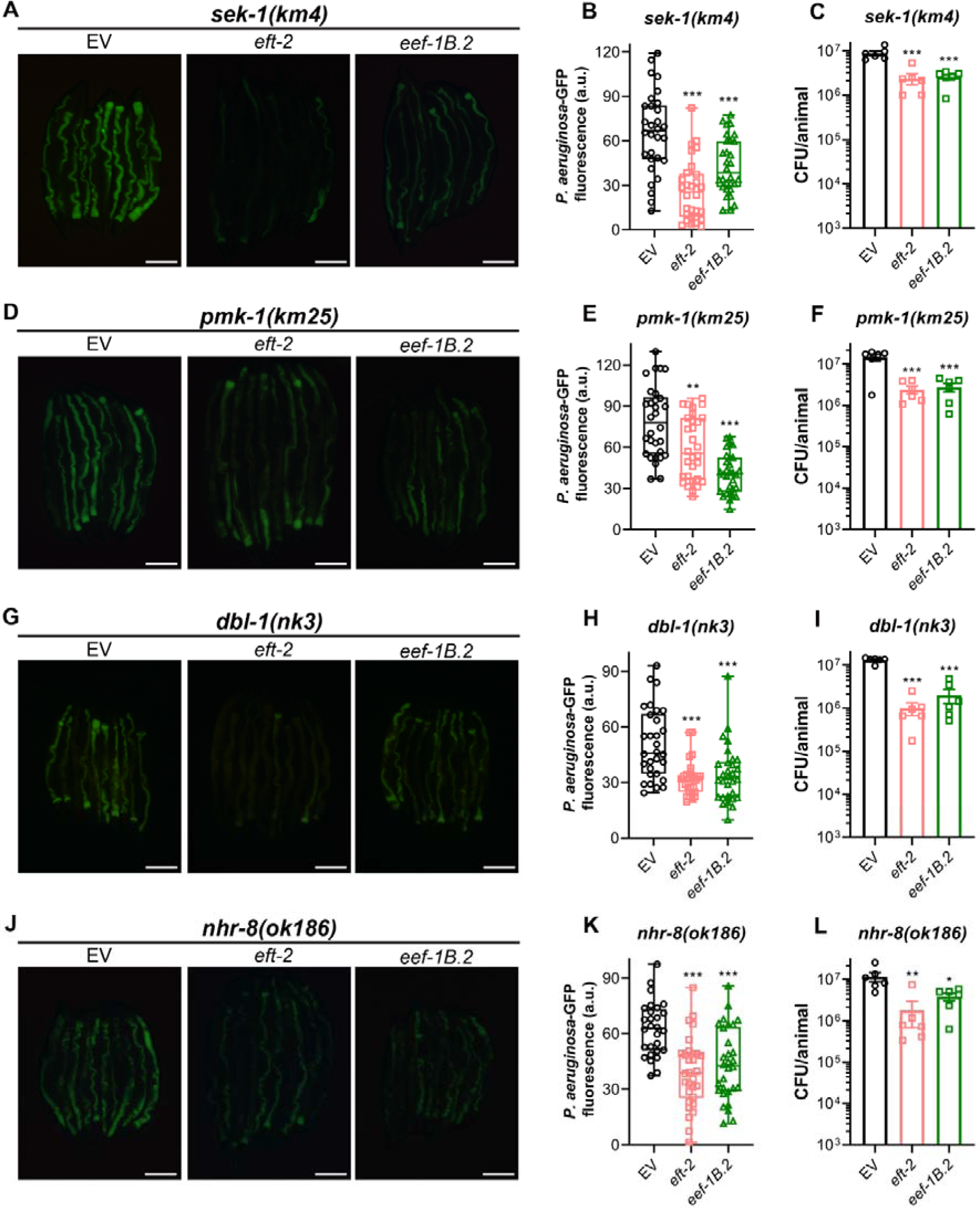
Inhibition of translation elongation reduces *P. aeruginosa* colonization in *sek-1(km4)*, *pmk-1(km25)*, *dbl-1(nk3)*, and *nhr-8(ok186)* animals. (A) Representative fluorescence images of *sek-1(km4)* animals incubated on *P. aeruginosa*-GFP for 12 hours at 25°C after treatment with the empty vector (EV) control, *eft-2*, and *eef-1B.2* RNAi. Scale bar = 200 μm. (B) Quantification of GFP levels of *sek-1(km4)* animals incubated on *P. aeruginosa*-GFP for 12 hours at 25°C after treatment with the EV control, *eft-2*, and *eef-1B.2* RNAi. \*\*\**P* < 0.001 via ordinary one-way ANOVA followed by Dunnett’s multiple comparisons test (*n* = 30 worms each). (C) Colony-forming units (CFU) per animal of *sek-1(km4)* animals incubated on *P. aeruginosa*-GFP for 12 hours at 25°C after treatment with the EV control, *eft-2*, and *eef-1B.2* RNAi. \*\*\**P* < 0.001 via ordinary one-way ANOVA followed by Dunnett’s multiple comparisons test (*n = 6* biological replicates). (D) Representative fluorescence images of *pmk-1(km25)* animals incubated on *P. aeruginosa*-GFP for 24 hours at 25°C after treatment with the EV control, *eft-2*, and *eef-1B.2* RNAi. Scale bar = 200 μm. (E) Quantification of GFP levels of *pmk-1(km25)* animals incubated on *P. aeruginosa*-GFP for 24 hours at 25°C after treatment with the EV control, *eft-2*, and *eef-1B.2* RNAi. \*\*\**P* < 0.001 and \*\**P* < 0.01 via ordinary one-way ANOVA followed by Dunnett’s multiple comparisons test (*n* = 30 worms each). (F) CFU per animal of *pmk-1(km25)* animals incubated on *P. aeruginosa*-GFP for 24 hours at 25°C after treatment with the EV control, *eft-2*, and *eef-1B.2* RNAi. \*\*\**P* < 0.001 via ordinary one-way ANOVA followed by Dunnett’s multiple comparisons test (*n = 6* biological replicates). (G) Representative fluorescence images of *dbl-1(nk3)* animals incubated on *P. aeruginosa*-GFP for 24 hours at 25°C after treatment with the EV control, *eft-2*, and *eef-1B.2* RNAi. Scale bar = 200 μm. (H) Quantification of GFP levels of *dbl-1(nk3)* animals incubated on *P. aeruginosa*-GFP for 24 hours at 25°C after treatment with the EV control, *eft-2*, and *eef-1B.2* RNAi. \*\*\**P* < 0.001 via ordinary one-way ANOVA followed by Dunnett’s multiple comparisons test (*n* = 30 worms each). (I) CFU per animal of *dbl-1(nk3)* animals incubated on *P. aeruginosa*-GFP for 24 hours at 25°C after treatment with the EV control, *eft-2*, and *eef-1B.2* RNAi. \*\*\**P* < 0.001 via ordinary one-way ANOVA followed by Dunnett’s multiple comparisons test (*n = 6* biological replicates). (J) Representative fluorescence images of *nhr-8(ok186)* animals incubated on *P. aeruginosa*-GFP for 24 hours at 25°C after treatment with the EV control, *eft-2*, and *eef-1B.2* RNAi. Scale bar = 200 μm. (K) Quantification of GFP levels of *nhr-8(ok186)* animals incubated on *P. aeruginosa*-GFP for 24 hours at 25°C after treatment with the EV control, *eft-2*, and *eef-1B.2* RNAi. \*\*\**P* < 0.001 via ordinary one-way ANOVA followed by Dunnett’s multiple comparisons test (*n* = 30 worms for *eft-2* and *eef-1B.2* RNAi, and 28 worms for EV control). (L) CFU per animal of *nhr-8(ok186)* animals incubated on *P. aeruginosa*-GFP for 24 hours at 25°C after treatment with the EV control, *eft-2*, and *eef-1B.2* RNAi. \*\**P* < 0.01 and \**P* < 0.05 via ordinary one-way ANOVA followed by Dunnett’s multiple comparisons test (*n = 6* biological replicates).

**Figure S10.**
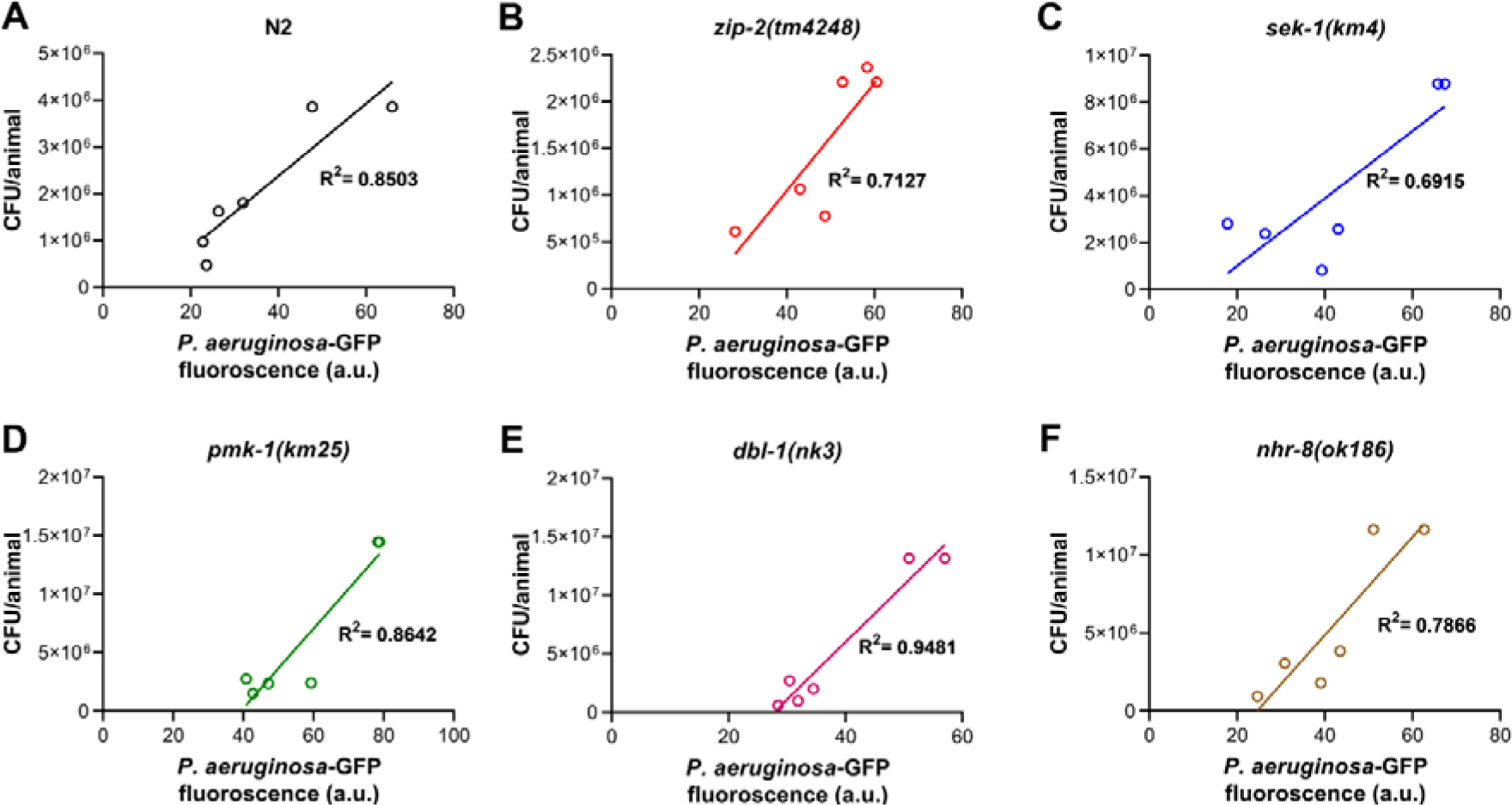
*C. elegans* gut *P. aeruginosa*-GFP levels correlate with colony-forming units (CFU) per animal. Correlation between *P. aeruginosa*-GFP levels per animal and CFU per animal for N2 (A), *zip-2(tm4248)* (B), *sek-1(km4)* (C), *pmk-1(km25)* (D), *dbl-1(nk3)* (E), and *nhr-8(ok186)* (F).

**Table S1:**
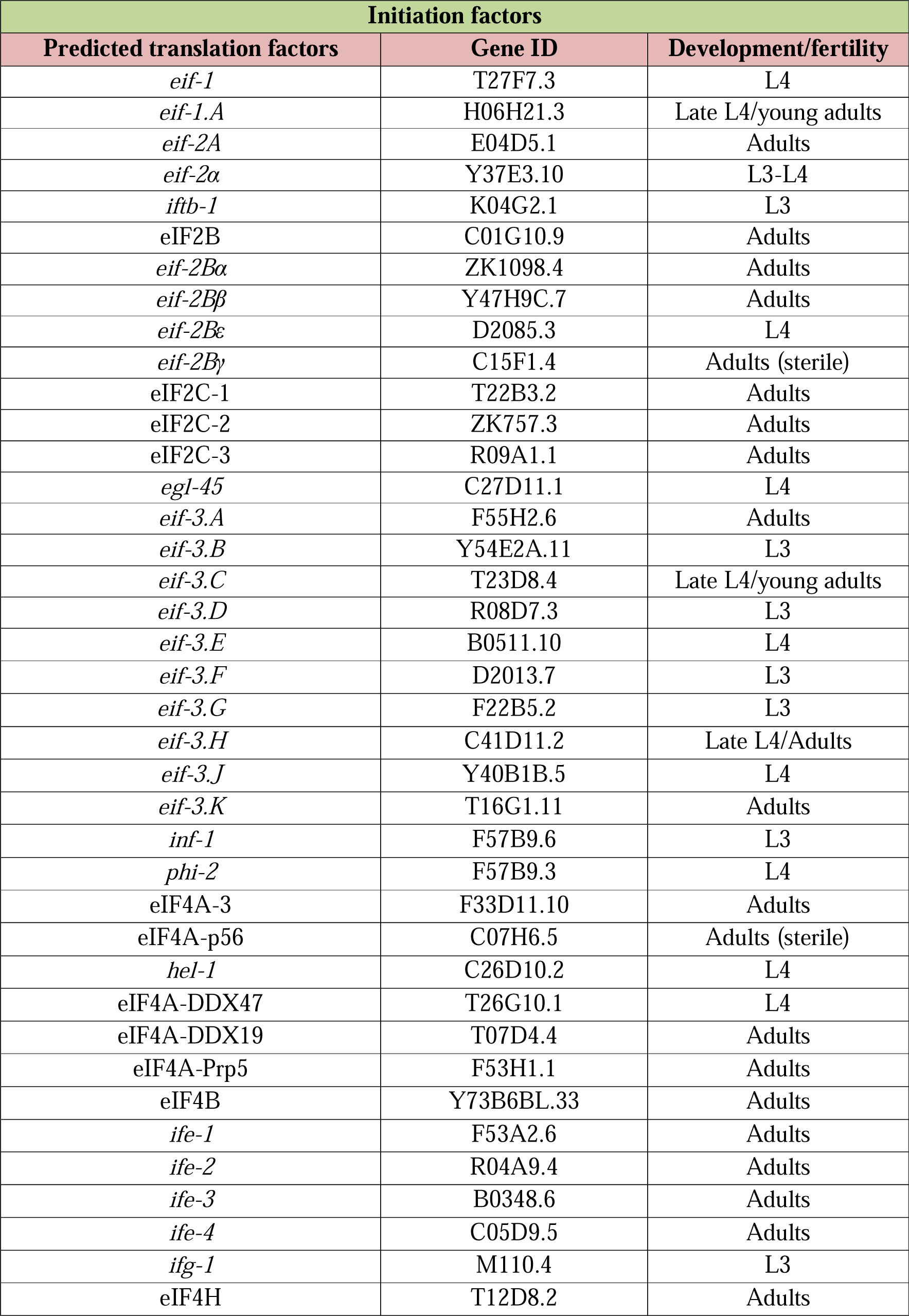
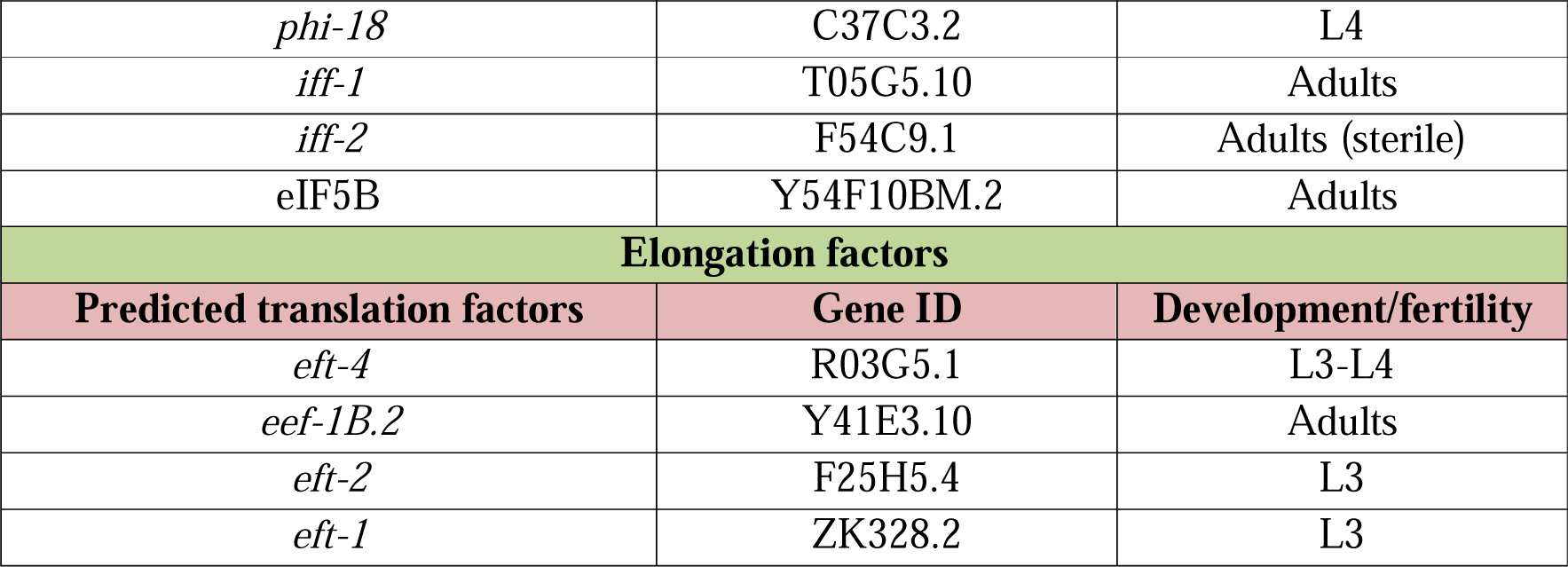
The list of *C. elegans* translation factors available in the Ahringer RNAi library and the effects of their knockdown on the development and fertility of N2 wild-type worms.

**Table S2:** Statistical analysis of survival curves from three independent experiments for each condition. A separate Excel file is provided.

**Table S3:** Upregulated and downregulated genes in *eif-2*α RNAi versus empty vector control RNAi in N2 animals. Genes exhibiting at least two-fold change and *P*-value <0.01 were considered differentially expressed. A separate Excel file is provided.

**Table S4:** Upregulated and downregulated genes in *eif-2*α RNAi versus empty vector control RNAi in *zip-2(tm4248)* animals. Genes exhibiting at least two-fold change and *P*-value <0.01 were considered differentially expressed. A separate Excel file is provided.

**Table S5:** Comparision of genes upregulated upon *eif-2*α RNAi in N2 and *zip-2(tm4248)* animals. A separate Excel file is provided.

**Table S6:** Upregulated and downregulated genes in *eft-2* RNAi versus empty vector control RNAi in N2 animals. Genes exhibiting at least two-fold change and *P*-value <0.01 were considered differentially expressed. A separate Excel file is provided.

**Table S7:** Upregulated and downregulated genes in *eft-2* RNAi versus empty vector control RNAi in *zip-2(tm4248)* animals. Genes exhibiting at least two-fold change and *P*-value <0.01 were considered differentially expressed. A separate Excel file is provided.

**Table S8:** Comparision of genes upregulated upon *eft-2* RNAi in N2 and *zip-2(tm4248)* animals. A separate Excel file is provided.

**Table S9:** Comparision of genes upregulated upon *eif-2*α RNAi and *eft-2* RNAi in N2 animals. A separate Excel file is provided.

**Table S10:**
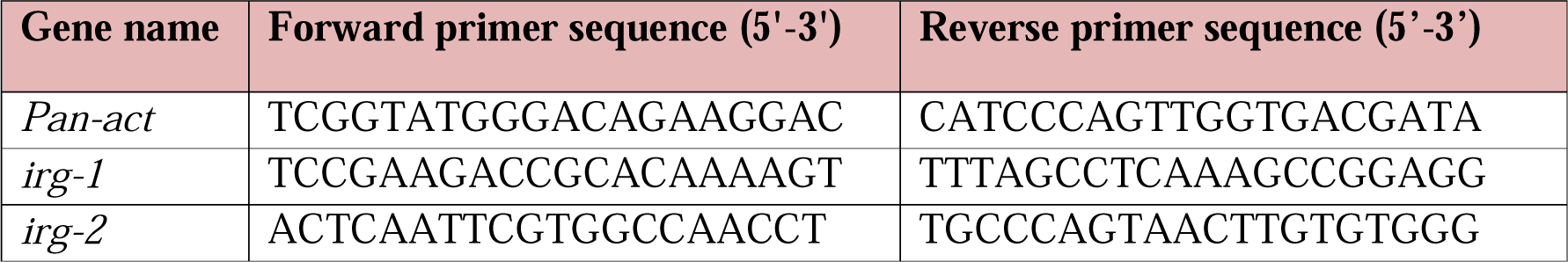
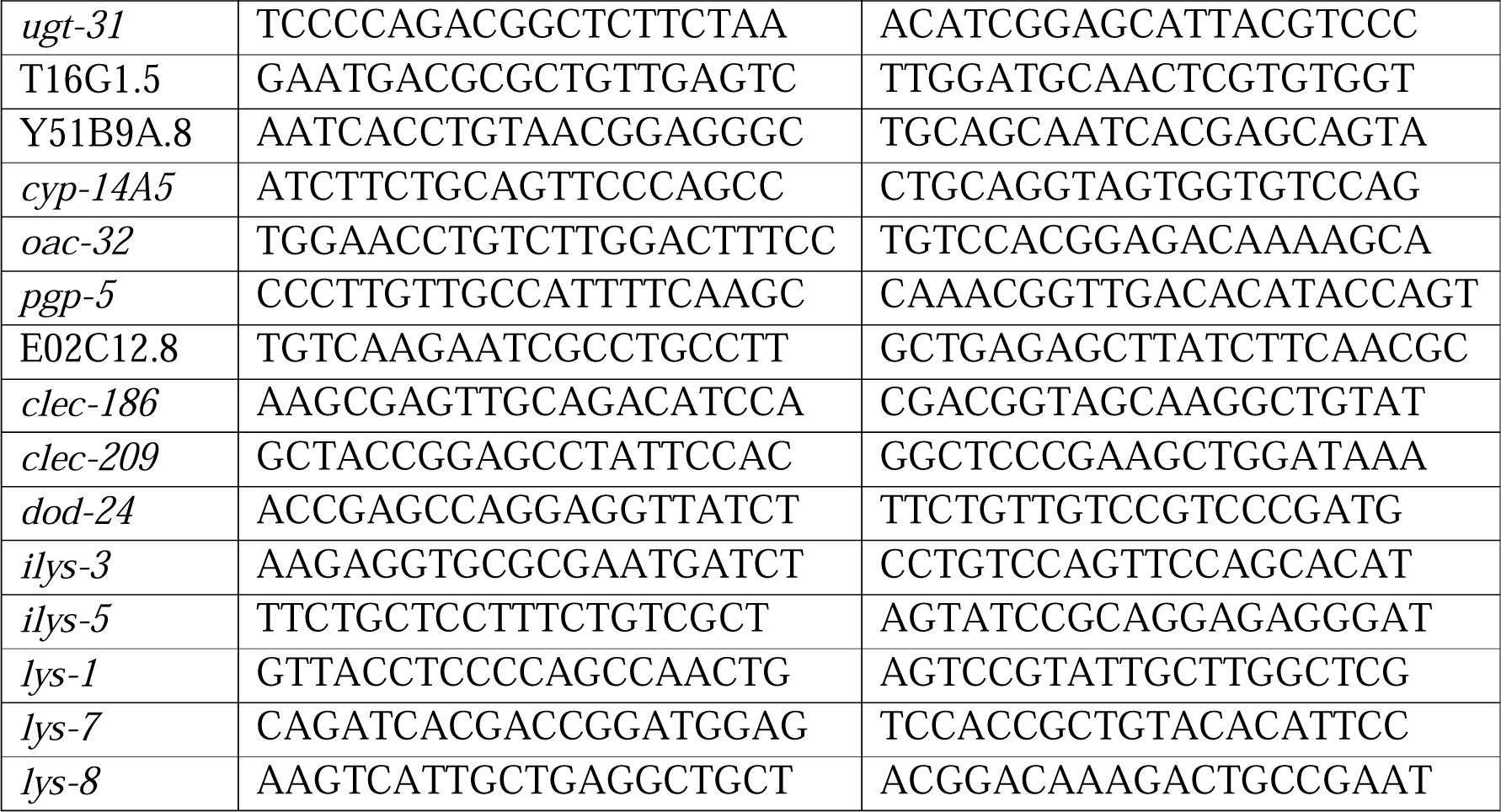
Quantitative reverse transcription-PCR primers used in the study.

